# Bee foraging preferences, microbiota and pathogens revealed by direct shotgun metagenomics of honey

**DOI:** 10.1101/2021.06.09.447678

**Authors:** Anastasios Galanis, Philippos Vardakas, Martin Reczko, Vaggelis Harokopos, Pantelis Hatzis, Efthimios M. C. Skoulakis, Georgos A. Pavlopoulos, Solenn Patalano

## Abstract

Honeybees (*Apis mellifera*) continue to succumb to human and environmental pressures despite their crucial role in providing essential ecosystem services. Owing to their foraging and honey production activities, honeybees form complex relationships with species across all domains, such as plants, viruses, bacteria (symbiotic and pathogenic), and other hive pests, making honey a valuable biomonitoring tool for assessing their ecological niche. Thus, the application of honey shotgun metagenomics (SM) has paved the way for a detailed description of the species honeybees interact with, in order to better assess the multiple factors governing their health. Here, we describe the implementation of optimized honey DNA extraction methodology coupled to direct shotgun metagenomics (Direct-SM) analysis, and to a computationally optimised and validated pipeline for taxonomic classification of species detected in honey. By comparing honey collected across 3 harvesting seasons in a stable apiary, we show that Direct-SM can describe the variability of sampled plant species, revealing honeybee behavioural adaptation. In addition, we reveal that Direct-SM can non-invasively capture the diversity of species comprising the core and non-core bacterial communities of the gut microbiome. Finally, we show that this methodology is applicable for the monitoring of pathogens and particularly for the biomonitoring varroa infestation. These results suggest that Direct-SM can accurately and comprehensively describe honeybee ecological niches and can be deployed to assess bee health in the field.

## 1. INTRODUCTION

Bees promote sustainable development goals by providing essential ecosystem services - most notably pollination - that collectively contribute to food security and maintenance of biodiversity (*1*, *2*). Yet, bees and, more visibly, managed honeybees, succumb to anthropogenic and environmental pressures, which synergistically increase the rate of colony loss (*3*–*6*). Over the past half century, pollination-dependent food production has tripled, while global honeybee stocks have increased by only about 45% (*7*). Consequently, honeybees are confronted with more intense beekeeping practices, increased hive density, and hive migration, which result in exposure and propagation of pathogens, such as *Varroa destructor* mites (*8*–*10*). In addition to these economic and social pressures, all pollinators are subject to various environmental stressors. Indeed, landscape alterations, caused by habitat loss and urbanisation, have dramatically reduced the abundance and diversity of food resources, leading to honeybee nutritional stress and the consequent weakening of their immune system, disruption of their gut microbiota, and increases in their susceptibility to viral and *Nosema* infections (*11*–*14*). Moreover, beehives providing pollination services are often placed in fields where they are exposed to varying degrees of herbicide and insecticide use. Exposure to non-lethal doses of pesticides alters the composition of bee microbial gut community, metabolic homeostasis and is likely to affect the gut microbiota-brain axis, with detrimental individual neurological effects, which collectively disrupt colony function (*6*, *15*–*19*). In parallel to these pressures on pollinators, the current climate crisis is likely to exacerbate their effects over the coming years (*20*–*23*). The challenge of building resilient food production systems while maintaining plant biodiversity and ecosystem stability necessitates the coordination of international biomonitoring strategies in order to comprehensively capture the multifactorial nature of threats to honeybee health (*24*, *25*).

A healthy honeybee colony results from a complex interaction between external factors (availability of food resources, climate, species interactions) and internal colony factors (honeybee physiology, degree of pathogenic infestation, behavioural and social stability) (*26*–*28*). Recently, several attempts have been made to develop colony health indices from various types of approaches (*29*, *30*). Among them, genomic technologies, such as DNA metabarcoding and metagenomics, are becoming attractive methodologies for tackling this challenge thanks to the constant increase in sequenced genomes and the increasing affordability of next generation sequencing (*31*–*33*).

These methods have already provided a wealth of information on seasonal plant exploitation and on honeybee adaptive behaviours with greater sensitivity and specificity compared to the standard analyses of pollen inspection by melissopalynology (*34*–*36*). For instance, recent studies have highlighted the frequent use of trees as a nutritional resource in urban and suburban environments, a clear preference for native species, and a capacity to diversify when preferred plants are no longer available (*34*, *36*–*39*). In addition, DNA metabarcoding of honeybee gut microbiota has revealed the conservation of 5 core members (*Gilliamella* sp., *Snodgrassella alvi*, *Lactobacillus* Firm-4 and Firm-5, and *Bifidobacterium* sp.), and the presence of additional non-core members (*Frischella perrara*, *Bartonella apis*) (*17*, *40*). Interestingly, recent studies also revealed individual-specific microbiomes, reflecting potential roles in adaptive functions, such as fluctuation of metabolic potential and nutritive resources across seasons, as well as social status (*41*–*43*). Contrary to plant exploitation and gut microbiome patterns, honeybee pathogens are rarely evaluated through genomic approaches. Instead, hive inspection is the standard method to assess infestation by *Varroa destructor* or brood diseases, while *Nosema ceranae* is detected mainly by conventional PCR, which often leads to delayed treatment initiation (*26*, *36*). Nevertheless, targeted approaches on certain pathogens revealed that co-infections could act synergically to disrupt gut bacterial composition (*44*, *45*). Notably, the specificity of certain viruses for targeting and interacting with core bacteria has uncovered their ability to modulate honeybee gut composition (*46*).

Such studies support the view that due to their extensive foraging activities, storage capacity and complex ecological interactions, honeybee colonies are unique large-scale biomonitoring tools that can provide insights into the status of ecosystems. Thus, it is imperative that, in order to implement effective biomonitoring strategies, a shift from targeted approaches towards practices able to quantify interactive pressures on honeybee colonies is needed.

Shotgun metagenomics on honey samples is an attractive methodology due to its non-invasiveness and potential for detecting many species interacting with honeybees. Honey is produced through the regurgitation (inversion) of flower nectar, which is subsequently placed in the comb until enough water evaporates. Throughout this procedure, the nectar (and, by extension, honey) comes in contact with a variety of organisms and therefore, contains DNA, termed environmental DNA (eDNA), from the sampled plants, the honeybee gut microbiome, and hive organisms, such as honeybees and others pathogens. Honey shotgun metagenomics has only recently been applied (*47*) and has been demonstrated to describe both the variety of the symbiotic and pathogenic organisms that honeybees come in contact with as well as the floral origin of honey (*48*, *49*).

However, this methodology requires further optimization, especially with regard to the deployment of inexpensive sampling methodologies and compatibility with fieldwork. In addition, an evaluation of bioinformatic pipelines, in particular taxonomic classification tools for the identification of honey-related species, is lacking. In practice, metagenomic classification tools are rarely optimized for particular applications, which may increase errors and skew results.

In the current study, we optimised a direct DNA extraction method to easily capture eDNA contained in honey and analysed the metagenomic composition by comparing taxonomic classifiers to analyse the landscape of species diversity contained in honey. By analysing samples collected over three different seasons in a static apiary, we demonstrated the great seasonal variability of plant exploitation by honeybees, which was in agreement with plant flowering periods and pollen analysis. Interestingly, we also detected the presence of all core and non-core gut bacterial populations at remarkably stable levels, underscoring the non-invasive nature of our approach for studying honeybee guts. Finally, we provide evidence that DNA levels detected by direct shotgun metagenomics can be used for the monitoring of *Varroa destructor* infestation, a major threat to colony survival. In conclusion, we have shown that our unique integrative biomonitoring method enables the simultaneous identification of the honeybee foraging behaviour, the state of gut microbial communities, and the presence of external pathogens, all of which have a strong influence on honeybee health.

## 2. METHODS

### 2.1. Apiary setup and monitoring

An apiary was installed on the property surrounding the BSRC ‘Alexander Fleming’ in Vari (Attika region, Greece) in December 2018 (GPS coordinates: 37°49’28.2” N 23°47’25.7” E). The colonies contained sister queens of the species *Apis mellifera macedonica*; all colonies started 9 months earlier. Hive populations and their degree of varroa infestation were monitored at least once a month throughout the entire year of 2019. Honeybee populations were estimated by measuring bee coverage on each side of the frames and scoring them from 1 to 10 (10 corresponding to maximum bee coverage with no space between the bees). The absolute population was calculated based on the estimation of complete frame coverage corresponding to 2000 honeybees (*50*). Varroa monitoring was performed using a wooden drawer installed at the bottom of each hive in order to monitor their natural falling off. To prevent fallen varroa from escaping or returning to the hive, this monitoring was optimised by covering the drawers with olive oil on baking paper. The degree of varroa infestation was then normalised per hive per day.

### 2.2. Honey collection and extraction

Honey was collected from four different hives in 2019 across 3 different seasons: spring (Hive 5), summer (Hives 6 and 7) and autumn (Hive 4). Collected frames from the hives were cut in small pieces (approximately 10-15 cm in length and 5 cm in width) and placed on top of a fine sieve on top of a glass bowl. The apparatus was placed in an incubator at 37°C for approximately 16 hours. This process allowed fresh honey (uncapped) to flow naturally. After 16 hours, the honey was collected and poured into clean glass jars, which were labelled and placed inside a drawer and kept at room temperature (RT: 18-24°C) until further processing.

### 2.3. DNA extraction

#### 2.3.1. Shotgun Metagenomics (SM) DNA extraction

DNA was isolated from honey from the 4 hives as in (*37*) with some modifications. For each hive, forty grams of honey were divided between two 50 mL Falcon tubes and filled with sterile distilled water up to 30 mL. Tubes were incubated in a water bath at 65°C for 30 mins, briefly shaken every 5 mins to ensure homogenisation, and ultra-centrifuged for 30 mins at 15,000 RPM using an SW50.2Ti rotor (Beckman Optima L-90K Ultracentrifuge). The supernatant was discarded and the pellets were pooled in 400 μL Buffer AP1 from the Qiagen DNeasy Plant Mini Kit (Qiagen). The mixture was homogenised progressively using the CAT X210 homogeniser for 40 seconds, avoiding the formation of foam. 80 μL of Proteinase K (1 mg/mL, Sigma) were added to the mixture and incubated for 50 mins at 65°C. During the incubation the tube was further inverted a few times every 15 mins. 4 μL of RNase A stock solution (100 mg/mL, from Qiagen DNeasy Plant Mini Kit) were added and the tube was briefly vortexed and incubated for 10 mins at 65°C. Further extraction proceeded according to manufacturer’s instructions (DNeasy Plant Mini Kit, Qiagen). with the following modifications: elution was performed with 25 μL of Buffer AE and DNA concentration was measured on a Nanodrop spectrophotometer. A total of 400 ng of DNA were sonicated in a total of 50 μL of Buffer AE. The solution was subsequently sonicated (Covaris S220 sonicator, temperature 7°C, 120 seconds) with the following parameters: duty cycle; 20%, Intensity 10; Cycle/burst; 10.

#### 2.3.2. Direct Shotgun Metagenomics (Direct-SM) DNA extraction

For each hive, five grams of honey were placed into a 15 mL Falcon tube. The tube was filled with sterile distilled water up to 10 mL and incubated in a hot water bath for 10 min, briefly shaken every 3 mins to ensure homogenisation. Several successive centrifugations at 14,000 g for 3 min in a microcentrifuge resulted in a single pellet in a 1.5 mL Eppendorf tube. The pellet was dissolved in 0.1 M NaOH, 5%Tween-20, vortexed for 30 seconds, incubated at RT for 15 min and quenched with 0.5 M Tris-HCl, 5 mM EDTA (. This is further referred to as the extraction mixture). The DNA was subsequently purified using Agencourt AMPure XP beads (Beckman Coulter), at a ratio of 2:1, according to manufacturer’s instructions. The purified DNA was snap frozen and stored at −80°C. Of note, the DNA for direct shotgun metagenomics was not sonicated.

### 2.4. Library preparation

A total of 8 DNA libraries (4 SM and 4 Direct-SM) were constructed using the Ion Plus Fragment Library Kit protocol (Thermo Fisher Scientific) with the following modifications: 5 to 10 ng of DNA were diluted, respectively, with sterile distilled water to a final volume of 39 μL. DNA was end-repaired by the addition of 10 μL of End-repair buffer and 1 μL enzyme per sample followed by 30min incubation at RT. Samples were purified using the AMPure XP beads (at a ratio of 1.9:1) and eluted in 20 μL Low TE. Adaptors were then ligated to the DNA in the presence of 5 μL ligase buffer and 1 μL ligase enzyme. 1 μL of universal IonXpressP1 adaptor was added in all samples with 1 μL of a barcoded IonXpress adaptor. The reaction was diluted in ddH2O to a final volume of 50 μL and incubated for 30 min at RT. After further purification with Agencourt AMPure XP beads (at a ratio of 1.5:1) and elution in 17.5 μL of Low TE, samples were amplified using 50 μL Platinum PCR Supermix High Fidelity and 2.5 μL Library amplification primer mix for 17 cycles (thermal cycling protocol: 72°C-20’/95°C-5’/ (97°C −15”,60°C-15”,70°C-1’)*17cycles/70°C-5’). A final 2 step purification was performed by adding 30 μL ddH2O to the 70 μL reaction, purified with AMPure XP beads (at a ratio of 0.8:1 to remove any fragments larger than 400 bp) and eluted in 20 μL ddH2O. The supernatant was used for a second purification using Agencourt AMPure XP beads (at a ratio of 0.5:1, total ratio 1.3:1 of initial). Library qualities and quantities were assessed on a Bioanalyzer using the DNA High Sensitivity Kit (Agilent Technologies). The quantified libraries were pooled at a final concentration of 7pM. The pools were processed, templated and enriched on an Ion Proton One Touch system. Templating was performed, using the IonPI^™^ Hi-Q^™^ OT2 200 Kit (Thermo Fisher Scientific) and sequencing with the IonPI^™^Hi-Q^™^Sequencing 200 Kit and the Ion Proton IonPI^™^V2 chips (Thermo Fisher Scientific) on a IonProton^™^ System, according to commercially available protocols.

### 2.5. Mock (simulated) samples

Mock samples were created to evaluate various steps and parameters of the computational pipeline. All mock datasets were created using FASTQsim (*51*) and a relative abundance (depth) was attributed to each species. Three mock samples were created, using the default IonTorrent parameters of FASTQsim, to evaluate the different taxonomic classification tools (see section below). The Viridiplantae community contained only Viridiplantae (11 plants and 1 green alga) while the non-Viridiplantae community contained 11 organisms: 4 non-Viridiplantae Eukaryota, 6 Bacteria, and 1 Virus. Honey community was created by merging the previous two mock samples.

### 2.6. Taxonomic classifiers and genomic aligners

The performance of five different computational tools for read taxonomic classification was evaluated. A brief description of the classifiers is described below.

#### 2.6.1. CCMetagen

CCMetagen is a metagenomic classification pipeline for a comprehensive and accurate identification of eukaryotes and prokaryotes in metagenomic samples. CCMetagen uses the KMA software for read mapping and alignment (*52*), which works in five steps: trimming of reads, heuristic *k*-mer mapping, fine alignment, ConClave scoring, and reference assembly. To enable high resolution of gaps and mismatches, KMA uses the Needleman-Wunsch alignment algorithm. ConClave alignment and scoring scheme allow a reference-guided assembly. CCMetagen ran using the pre-indexed nt database from 2018 (downloaded from: https://researchdata.ands.org.au/indexed-reference-databases-kma-ccmetagen/1371207)

#### 2.6.2. DIAMOND

DIAMOND is an application to align a set of sequences, which can be used as a query against a sequence database of preference (e.g., nr-NCBI) (*53*). DIAMOND follows a seed-and-extend approach and uses a double indexing (both for queried sequences and background database) for better performance. The latest versions give comparable results to BLAST at a much faster speed and have become a widely used application in the field of metagenomics.

#### 2.6.3. Kraken2

Kraken2 is an exact *k*-mer-based approach for accurately and quickly classifying metagenomic reads (*54*). Compared to its predecessor, Kraken2 comes with advanced indexers and data structures for faster database building and querying; thus, it offers high-quality analyses and fast classification speeds. The main idea behind Kraken2 is to first match each *k*-mer within a query sequence to the lowest common ancestor (LCA) of all genomes containing the given *k*-mer. In this study, Kraken2 was used with its default parameters with an updated version of the nt database (May 2020).

#### 2.6.4. MG-RAST

MG-RAST is an online service for automatic phylogenetic and functional analysis of metagenomes and is accessible through https://www.mg-rast.org. It is a pipeline consisting of many specialized tools for quality control, classification, annotation, visualization and functional annotation of reads (*55*). MG-RAST is one of the most widely-used services in the field of metagenomics, as one can upload raw data in fastq format and compare them against 458,163 metagenomes containing 1,922 billion sequences (stats 05/21).

#### 2.6.5. Minimap2

Minimap2 is an alignment application to map DNA or long mRNA sequences against a large reference database and is recommended for shorter reads (*56*). For ~10 kb noisy read sequences, minimap2 was found to be faster than mainstream long-read mappers, whereas for >100 bp short reads, minimap2 was found to be three times faster than BWA-MEM and Bowtie2.

### 2.7. Filtering and normalisation of the libraries

#### 2.7.1. Sequence pre-processing

Adaptors and low-quality reads were automatically removed after having been sequenced by the IonTorrent Server. Samples were split according to barcode.

#### 2.7.2. Filters based on Kraken 2 confidence score and usual laboratory cross-contaminant species

To address whether the taxonomic classification is influenced by the number of species in each mock sample, by the library size, or by the fragment length, additional mock samples were tested to decide which Kraken 2 confidence score should be applied. In kraken2, a perfect hit will have all the *k*-mers mapped. In case of mismatches, all *k*-mers that contain this nucleotide will be impacted. The confidence score represents the number of matching *k*-mers divided by the total number of *k*-mers. The confidence thresholds were evaluated using three metrics:

i. Number of species: Calculated as the total number of species.
ii. Root-mean-square-error:

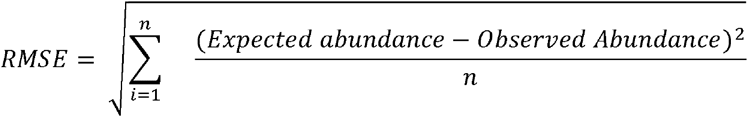

where *Expected abundance* is the set abundance in the simulated sample and the *Observed abundance* is the reported abundance from kraken2 at the tested confidence threshold. The RMSE was used because it evaluates the overall error in the classification and abundance estimation.
iii. Sensitivity:

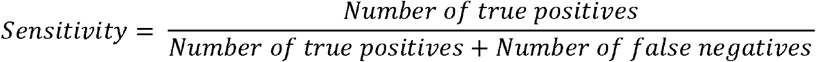

where the *Number of true positives* is the number of species in the mock that were correctly detected and the *Number of false negatives* is the number of species in the mock that were not detected (reported as 0 reads in kraken2). Sensitivity is the ability to correctly identify the species that do exist in the sample.
iv. Positive predictive value:

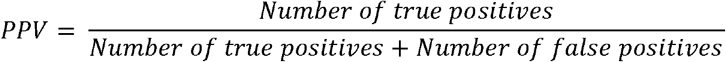

where the *Number of true positives* is the number of species in the mock that were correctly detected and the *Number of false positives* is the number of species reported by kraken2 but not present in the mock.

The filtered kraken2 files generated after the processing of the 8 sequenced libraries were imported and processed with the *taxonomizr* R package to obtain the taxonomic attribution for each species. Reads assigned to Phylum Chordata were removed along with the family *Drosophilidae*, the genus *Drosophila*, and the species *Drosophila melanogaster* due to possible laboratory cross-contamination.

#### 2.7.3. Read normalisation across libraries

DESeq2 version v3.11 was accessed via the Bioconductor library and read normalization was performed with the use of the *estimateSizeFactors* function, which applies the relative-log expression (RLE) (*57*). Notably, DESeq2 RLE transformation has been shown to be appropriate for normalisation of shotgun metagenomic libraries (*58*).

### 2.8. Statistical analysis and R packages

The analysis and graphical visualisation of the results were performed using R versions 3.6.3 and 4.0.3 on a x64 system running Microsoft Windows 10 Pro. Statistical analysis was performed in R and Prism9.

#### 2.8.1. DESeq2 differential analysis and clustering

Following library normalisation, DESeq2 was used for differential abundance analysis. First, the methods were used as ‘contrast’ to investigate which species differ significantly between methodologies. Second, the likelihood-ratio test (LRT) was used to identify the species that differ significantly across seasons. In order to avoid aberrant contributions of low-coverage species, additional filtering of the species identified by fewer than 50 reads between two libraries (to maintain the Direct-SM and SM pairing) was applied for hierarchical clustering. Then, using the DEGreport package (*59*), we identified the clusters of species whose abundance differed similarly across seasons. This was achieved using the degPatterns function with the minimum number of species per cluster (minc parameter) set at 2.

#### 2.8.2. Principal Component Analysis (PCA)

PCA was performed across normalised libraries quantification and filtered for species with fewer than 50 reads coverage. PCA was performed using the prcomp function (base R) and visualisation was performed using the factoextra package (*60*).

### 2.9. Plant species validation

The plants which were detected and identified, were validated by various analyses. First, their presence was validated using the BIEN database (*61*) and by comparing their occurrence within defined geographic boundaries across the world. Second, a gyroscopic analysis was performed by the CheMa laboratory on each honey sample (https://www.chema.gr, Corinthos, Greece). The occurrence and abundance of each pollen family were then compared with families identified by metagenomics after Kraken2 classification and DESeq2 normalisation at this level (with a min of abundance of 50 reads for each family). Third, a visual inspection and botanical characterisation was undertaken within the area using an online database of Greek plants (https://www.greekflora.gr), in which we also extracted the flowering pattern of species present in Greece.

### 2.10. Literature search to assess the relationship of non-plant species with honeybees

In order to attribute a relationship between honeybees and the detected non-plant species, we searched for PubMed articles which contained the words ‘bee’/’honey’/’apis’ in the title and/or abstract’ and which were associated with the species name. We limited our analysis to the species initially detected with a coverage higher than 100 reads. NCBI abstract hits were manually inspected to confirm the relationship of each species with honeybees. 6 relationships were attributed: “Bacterial gut community”, “Host”, “Human cross contamination”, “Others”, “Pathogen”, “Unknown”.

### 2.11. Functional annotation

Reads belonging to the bacterial species showing significant increase during the autumn season were extracted using a script from KrakenTools (https://ccb.jhu.edu/software/krakentools/index.shtml). The reads were aligned to a pre-indexed Uniref90 database using DIAMOND (*53*) blastx with the following parameters: --mid-sensitive -b8 -c1. The resulting table was regrouped and renamed to Gene Ontology (GO) annotation using scripts from the HUMAN 3 pipeline (*62*). Normalised frequencies of GO attribution were then analysed to identify the most enriched term for each species.

### 2.12 Genomic re-alignment of sequenced libraries against the varroa genome, calculation of the means of natural varroa fall and correlation with metagenomic data

For validation purposes, the reads from each library (fastq format) were re-aligned to the *Varroa destructor* genome using HISAT2 (*63*), with the following parameters: -k 1 -p 12 --mp 30,30 --rdg 30,30 --rfg 30,30 −dta. Aligned reads were normalised with total sequenced reads (counts per million) and correlated with the average of the natural fall of varroa/day over periods of 2 weeks, 1 month and 2 months and with the respective abundances of *Varroa* species attributed by the metagenomic pipeline.

## 3. RESULTS

### 3.1. Experimental design and description of the direct shotgun metagenomic honey DNA extraction method

Our study took place in an apiary whose ecosystem in a 2 km radius (12.5 km^2^), is typical of a coastal, semi-arid Mediterranean climate and is mainly composed of conifers, pines, dense shrubs, and olive trees (31%), as well as urban areas containing many ornamental gardens (59%), coastline, and sea area (10%). It is located on the edge of the Hymettus Mountain on the suburbs of Athens in Greece (Figure 1a). The four hives of this study showed comparable evolution of honeybee population growth and varroa infestation at the time of honey collection (Sup. figure 1).

**Figure 1:**
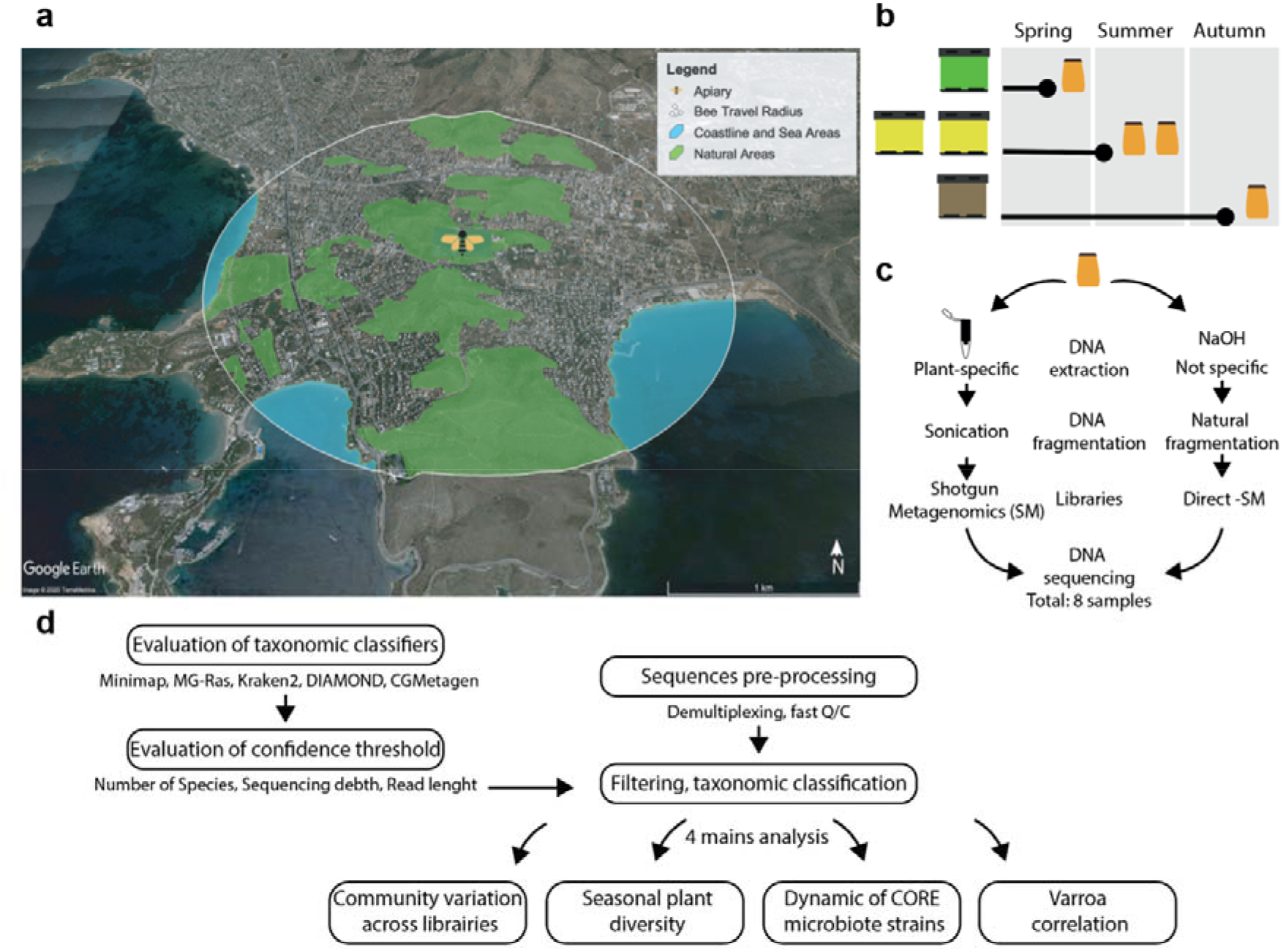
Overview of the experimental design and bioinformatic workflow. (a) Apiary location and habitat structure. (b) Honey sampling across 3 seasons. (c) 8 libraries were sequenced following 2 methodologies: Shotgun Metagenomic (SM) and direct-SM library preparation. (d) Flowcharts depicting the detailed bioinformatics workflow leading to the four main output analyses.

A total of four honey collections were carried out, representative of the main beekeeper harvest seasons, in spring, summer, and autumn (Figure 1b). For each harvest, two DNA extraction techniques were compared (Figure 1c). The first technique follows a classic metagenomic shotgun (SM) protocol for extracting DNA from 40 g of honey using specific extraction columns, followed by a sonication step to fragment the DNA before the preparation of a sequencing library (*37*, *47*). The second technique is more direct (Direct-SM) and has been developed using 5 g of honey. The lysis and DNA extraction does not involve any specific purification kit, but instead uses an optimised NaOH-based buffer (See methods). The DNA contained in honey is already fragmented (Sup. figure 2a); therefore, Direct-SM involves no sonification.

A total of 8 libraries were prepared and sequenced (4 replicates for the Direct-SM and 4 replicates for the SM), (Sup. figure 2a) and analysed according to specific bioinformatic workflows, including the initial evaluation of various taxonomic classifiers (Figure 1d).

### 3.2. Kraken2 performs best for metagenomic classification of simulated honey samples

In order to choose the most appropriate bioinformatic workflow for the analysis of shotgun metagenomics data from our honey samples, we first evaluated several taxonomic classifiers using mock (simulated) samples of various species communities. The primary reason to apply shotgun metagenomics in honey samples was to simultaneously study the foraging behaviour and health of *Apis mellifera*. Therefore, we built 3 different communities of mock samples. The first included 12 known pollinated plant species (Viridiplantae community). The second included 11 non-plant eukaryota, bacteria and viruses, such as *Apis mellifera*, the putative pathogens *Varroa destructor*, *Apis mellifera filamentous virus* and *Nosema ceranae*, and some symbiotic bacteria (non-Viridiplantae community). Finally, we also built a simulated honey sample containing all 23 species from both Viridiplantae and non-Viridiplantae communities (Honey community) (Sup. figure 3a).

We assessed several taxonomic classification tools based on their ability to quantitatively and correctly assign the mock sequence reads distribution at both genus and species levels. Genus distribution of the Viridiplantae community shows that the DIAMOND classifier followed by the kraken2 classifier, produce the closest distributions to the mock (Sup. Figure 3b). In contrast, when these classifiers were evaluated for taxonomic attribution at the species level, kraken2 performed better than DIAMOND for both the Viridiplantae and non-Viridiplantae communities (Sup. Figure 3c-d). Finally, when using the simulated honey community, kraken2 again produced the greatest correlation and significance between expected abundances and observed abundances at the species level (Figure 2). The reliability performance of Kraken2 in identifying taxa at the species level was further confirmed by the calculation of the root mean square error (RMSE) (Sup. Figure 3e), so this taxonomic classifier was chosen for the remaining analyses.

**Figure 2:**
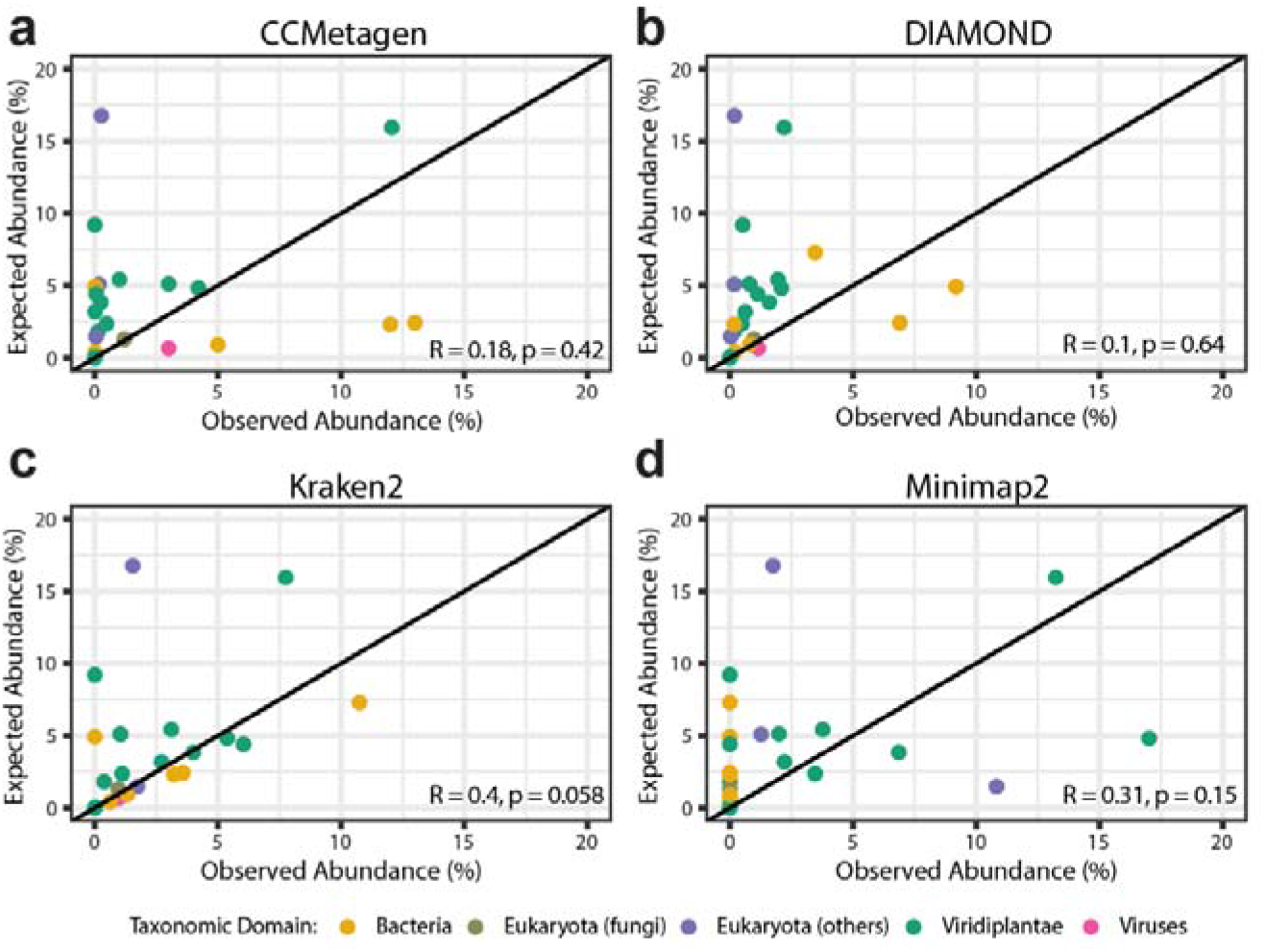
Kraken2 provides optimal performance for taxonomic classification and genomic alignment. Correlation between expected and observed abundance from the mock honey species community using CCMetagen (a), DIAMOND (b), kraken2 (c) and minimap2 (d).

Since kraken2 at default settings was found to report a much higher number of species than those contained in the simulated honey community (above 5000, Sup. figure S4a), we sought to determine the highest confidence threshold for this classifier. For this, we tested several scenarios likely to influence classification: The initial number of species contained in a simulated sample (23 versus 64 species, Sup. figure 4), the size variability of our sequencing libraries (1 versus 8 million reads, Sup. figure 5), and the influence of the length of the sequenced fragments (110 versus 250 bp, Sup. figure 6). These analyses showed that the absolute number of assigned species drops significantly upon application of a confidence level of 0.1 and approaches the actual composition at around 0.5 (approximately 180 species reported) in every scenario. In addition, our analysis revealed greater robustness of classification when samples contained higher numbers of species and more sequencing reads, while the length of the fragments did not seem to influence the classification. On the other hand, the sensitivity dropped significantly after the 0.5 threshold in most scenarios. Therefore, kraken2 with a confidence threshold of 0.5 was selected for analysis of the experimental honey samples as it shows the minimal errors, the highest sensitivity and limited lost reads.

### 3.3. Direct-shotgun metagenomic analysis reports similar species distribution and abundance compared with standard shotgun metagenomic analysis

Between 1.4 and 8 million reads were obtained after sequencing of the 8 honey samples (Table 1). After taxonomic classification with kraken2 with a cut-off of 0.5 and additional filtering for putative laboratory cross contaminants, 50,000 to 600,000 reads per library were available for downstream analyses. On average, 85% of the reads were classified at the species level, 9% at the genus level, and fewer than 4% at the family level (Table 2).

**Table 1.** Distribution of raw, classified and filtered sequencing reads across the 8 libraries

| Sample | Raw reads | Kraken2 Classified reads | Classified (%) | After filtering (Confidence 0.5) | After filtering (% of classified) |
| --- | --- | --- | --- | --- | --- |
| DirectSM_H4 | 3,282,781 | 480,566 | 14.6% | 230,316 | 47.9% |
| SM_H4 | 4,058,520 | 730,745 | 18 % | 440,017 | 60.2% |
| DirectSM_H5 | 1,444,924 | 154,577 | 10.7% | 58,722 | 38% |
| SM_H5 | 8,160,732 | 253,515 | 3.1% | 143,154 | 56.5% |
| DirectSM_H6 | 2,379,244 | 676,415 | 28.4% | 274,230 | 40.5% |
| SM_H6 | 3,619,844 | 1,247,102 | 34.4% | 623,454 | 50% |
| DirectSM_H7 | 2,280,627 | 827,329 | 36.3% | 482,892 | 58.4% |
| SM_H7 | 2,038,988 | 1,353,429 | 66.3% | 223,023 | 16.%% |

**Table 2.** Kraken2 taxonomic classification across the 8 libraries

| **Sample** | **Family (%)** | **Genus (%)** | **Species (%)** |
| --- | --- | --- | --- |
| DirectSM_H4 | 15415 **(6.69%)** | 40710 **(17.68%)** | 174190 **(75.63%)** |
| SM_H4 | 22336 **(5.08%)** | 91293 **(20.75%)** | 326388 **(74.18%)** |
| DirectSM_H5 | 4686 **(7.98%)** | 8966 **(15.27%)** | 45070 **(76.75%)** |
| SM_H5 | 6587 **(4.6%)** | 12582 **(8.79%)** | 123985 **(86.61%)** |
| DirectSM_H6 | 4318 **(1.57%)** | 10269 **(3.74%)** | 259643 **(94.68%)** |
| SM_H6 | 6056 **(0.79%)** | 20188 **(3.24%)** | 597210 **(95.79%)** |
| DirectSM_H7 | 5082 **(1.05%)** | 3585 **(0.74%)** | 474225 **(98.21%)** |
| SM_H7 | 1920 **(0.86%)** | 1020 **(0.46%)** | 220083 **(98.68%)** |

The number of species identified in the 8 libraries showed great variation, from 265 to 663 species (Figure 3a). However, once species were categorised into domains, we observed a remarkable conservation of their distribution. Indeed, the majority of identified species are classified under the domain of Bacteria (53 ± 6.7%), then Viridiplantae (31 ± 6%), Eukaryota (14 ± 2.4%) and a minority of viruses (2.1 ± 0.5%). No significant differences in distribution were detected when the two techniques were compared (SM versus Direct-SM, two-way ANOVA, p = 0.2795) or when the domains were compared individually (SM versus Direct-SM, Sidak’s Multiple comparison tests, Adj. p > 0.3). In addition, more than 75% of the identified species overlap between the 2 techniques and for the 4 analysed hives (Figure 3b), pointing to a limited influence of DNA extraction methodology to species identification.

**Figure 3:**
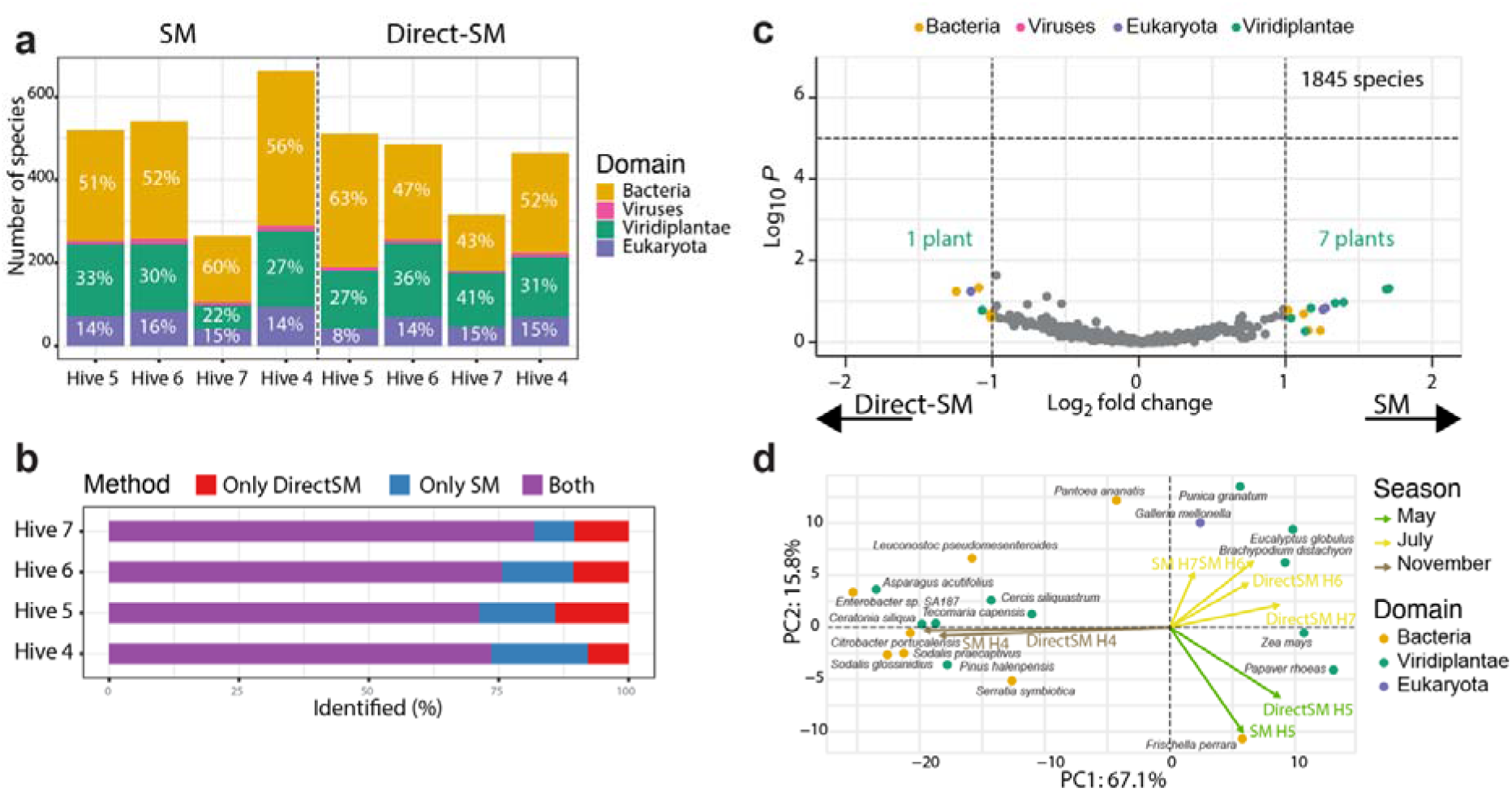
Seasonal variation, not extraction methodologies, explain variations between libraries. (a) Barplot of the distribution of the number of identified species per library and classified according to the domain they belong to. (b) Barplot of percentages of overlapping species between SM and Direct-SM methodologies for each hive. (c) Volcano plots of species abundance associated with SM and Direct-SM methodologies. Dashed lines indicated the limit of significance (y axis: Walt test with BH multiple comparison, x axis: log2 fold change −1 < or > 1). (d) PCA analysis across identified species. Arrows correspond to the 8 samplings across seasons and each dot corresponds to the species that contributed the most to the seasonal deviation.

Initially, the sequenced data were quantitatively normalised over all libraries using DESeq2 (see Methods) and compared at both family and species levels. The comparison between the two methodologies did not identify any significant abundance variations at any level (DESeq2 Wald test, corrected for multiple testing using BH). However, 19 families (3.9%) and 21 species (1.1%) were respectively detected with a log2-fold change greater or less than 1, with a slight deviation in abundance towards Viridiplantae when the DNA purification technique used was plant-specific (SM) (Figure 3c, Sup. figure 7). Even if these observations confirm a slight bias introduced by the SM extraction method, overall, they suggest a very weak impact of the methodology for the determination of the metagenomic content and abundance of DNA in honey.

To further verify the consistency between the two methodologies, we performed hierarchical clustering and principal component analyses. Both revealed a clustering of the samples largely driven by seasonal changes. Despite our modest sample size, this seasonal grouping is explained by the consistency of determination of species abundance between the SM and Direct-SM methodologies, but also by the identification of clusters comprising, as expected, season-specific plant species and seasonal bacterial, eukaryotic, and viral loads (Figure 3d, Sup. figure 8). For instance, the wax moth, *Galleria mellonella*, was consistently found in samples extracted during summer, when honeybee colonies are more prone to infestation by this pathogen (*64*). These analyses demonstrate that Direct-SM has the potential to simultaneously describe the honeybee foraging pattern and the dynamics of honeybee interactions with other species.

### 3.4. Metagenomic analyses accurately describe seasonal plant species variability

Normalised but unfiltered sequencing data reveal that 398 species belonging to Streptophyta were potentially foraged around the apiary. Despite this large diversity, only 45 genomes were covered by at least 50 reads, which correspond to 25 families and 41 genera (Sup. table 1). To verify how accurate the identification of the main plant species was, we used three types of validation: 1) *in silico*, by comparing our samples with the largest publicly accessible plant databases from the Botanical and Ecological Information Network (BIEN), 2) by melissopalynology (pollen inspection) on the 4 collected honey samples and 3) *in situ*, by examination of plant species in the area.

Using the BIEN database, we show that 66 ± 5.02% of our plant species overlap with species identified within the area of the Mediterranean basin, with the greatest overlap in Spain (Sup. figure 9). As an additional validation step, we repeat this analysis using an equivalent area along the same latitude for Asia and for North America. As expected, significantly lower overlap was observed in Asia (51 ± 3.88%); however, a significant overlap (85 ± 1.65%) was observed for North America highlighting, among other things, the overwhelming preponderance of north-american plants in the BIEN database.

Pollen identification in honey can rarely discern plant origin at the species level and, therefore, we restricted comparison with metagenomic data to family level only: we found that 11 out of 20 families identified by pollen analysis were also present in our metagenomic analysis with great seasonal consistency ranging from 50% to 87.5% of overlap (Sup. figure 10a-b). Interestingly, this comparison revealed foraging patterns not necessarily associated with pollen collection, such as visits of Arecaceae, Pinaceae, and Vitaceae families (Sup. table 2). Conversely, several families abundantly present in pollen analysis were lacking from the metagenomic analysis, highlighting the lack of sequenced genomes belonging to the Araliaceae or Boraginaceae families. Finally, correlations between abundance of families identified by both metagenomic methods and pollen analysis were significant only for 2 of the 4 collected honey samples, highlighting the limitations of pollen analysis to quantitatively assess the botanical origin of honey (Sup. figure 10c).

The most robust approach to validate the plants identified through shotgun metagenomics was implemented using a local database of Greek plants coupled with visual inspection of the area. Indeed, among the 45 species identified by our bioinformatic pipeline, 20 were correctly assessed at species level, 18 at genus level and only 7 hits were considered false positives (Figure 4a, Sup. table 1). The overall abundance of plant species and genera found in honey samples were in agreement with the environment surrounding the apiary, which mainly comprises trees: *Pinus* (pine tree, 47.42%), *Phoenix* (Date palm tree, 12.5%), *Eucalyptus* (Eucalyptus tree, 12.48%) were the most represented (Figure 4a). Finally, the dynamic analysis of plant abundance across seasons revealed that 16 species were significantly enriched in honey samples harvested in a specific season, such as *Papaver* (poppy, 13.98%) during the spring season, and that this enrichment overlapped or closely juxtaposed the flowering period of these plants (Figure 4b-c), thus reinforcing the validity of our analysis.

**Figure 4:**
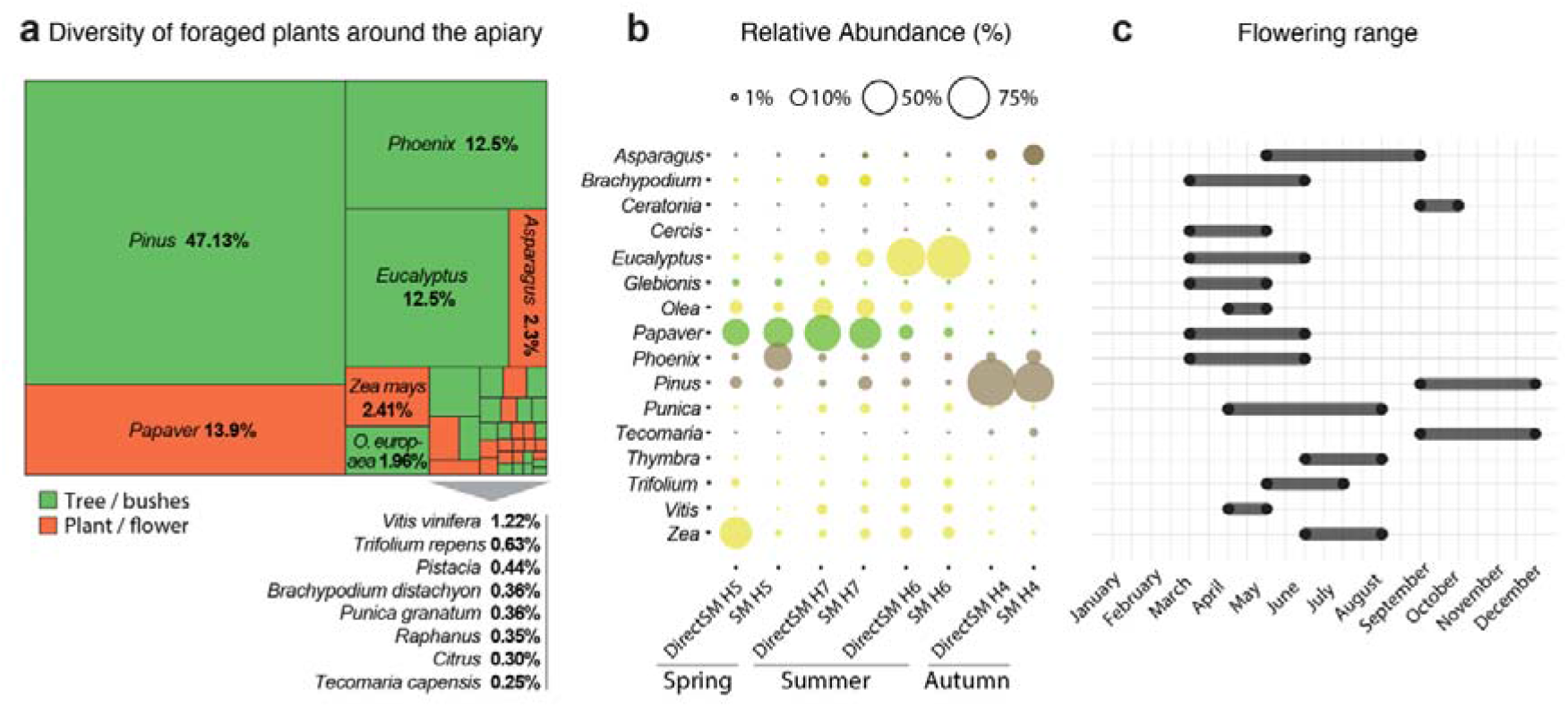
Plant distribution and abundance capture composition and seasonal changes of apiary area. (a) Abundance treemap of the 38 validated plants. Plants validated at the species level are presented in full, others by genus only. The % indicates the general abundance across all honey samples (only species with abundance higher than 0.2% are indicated). (b) Dynamics of abundance across hives and seasons. The colour code highlights significant direction toward a specific season (spring: green, summer: yellow, autumn: brown). Seasonal significance was obtained with DESeq2 using a likelihood-ratio test (see Methods). (c) Flowering ranges of the plants shown in panel (a).

Overall, and despite the lack of local plant genomic data, these analyses show that our metagenomic pipeline accurately describes the botanical origin of honey samples and their seasonal variability with a greater accuracy than *in silico* latitude occurrence or pollen analyses.

### 3.5. Non-invasive characterisation of the hive pathogens and the dynamics of core and non-core honeybee gut microbiome

In line with previous work (*47*, *49*), our pipeline revealed that the majority of DNA traces contained in honey samples did not belong to plants (Figure 3a). To better understand the importance of this diversity and their relationship with honeybees, we first used a text mining approach to categorize their relationship with honeybees based on current literature (See method and Sup. table 3). After filtering out reads emanating from the *Apis mellifera* filamentous virus, whose DNA is represented in more than 86% of all sequenced reads (Sup. figure 11a), we found that 75% of non-plant species were constituents of the gut bacterial community of honeybees, dominated by *Lactobacillus kunkeii* (Figure 5a); 4.6% originated from bacteria commonly present on human skin—likely due to beekeeper interventions inside the hives—and only 1.1% were described as pathogenic for honeybees, including traces of the seasonal opportunistic bacterium *Spiroplasma melliferum* (*65*) (Sup. figure 11b). Interestingly, 6.8% of DNA present in honey samples had no direct relation to honeybees, but mainly arose from 2 plant pathogens and an aphid symbiont (Sup. table 3).

**Figure 5:**
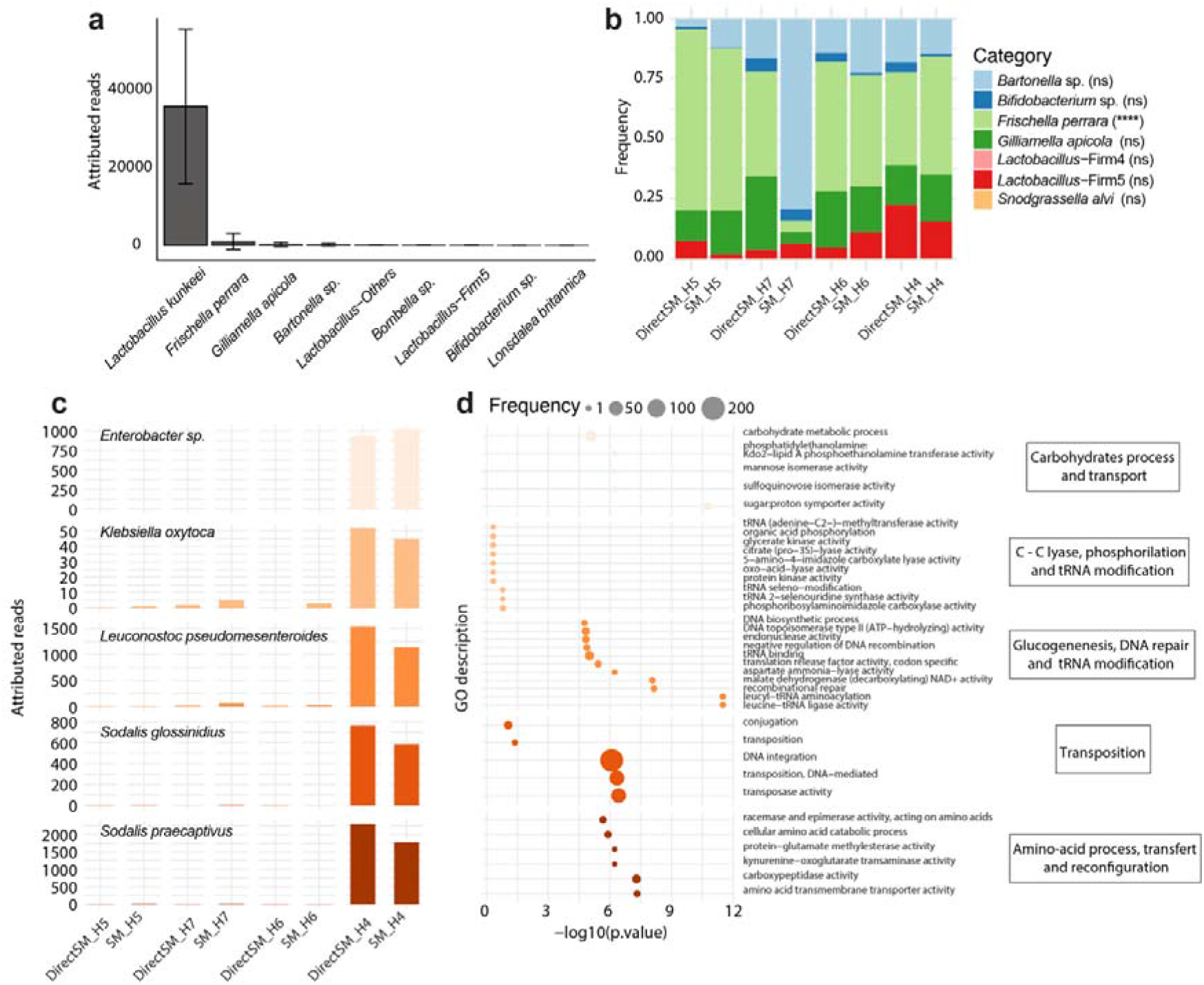
Characterisation of core honeybee microbiota stability and seasonal dynamics of five non-core bacterial strains. (a) Distribution of the 10 most abundant microbiota species. (b) Barplots of symbiotic bacteria distribution of species previously described as core or non-core. Species with significant variations are annotated with stars (*L. kunkeei* was excluded). (c) Distribution of the five species specifically associated with autumn honey samples with (d) their respective gene functional enrichment categories. Only the most representative terms among the top 20 significant GO terms are shown.

We then characterised the dynamics of the gut community and found remarkable conservation of abundance in the harvested honey samples, independently of season (Figure 5b). All previously described core members, such as *Bifidobacterium* sp. (2.6 ± SEM 0.7%), *Gilliamella apicola* (18.1 ± SEM 2.7%), *Snodgrassella alvi* (< 0.1%), *Lactobacillus* Firm-4 (< 0.1%), Firm-5 (9.1 ± SEM 2.4%), and non-core members *Frischella perrara* (47.5 ± SEM 7.5%), *Bartonella apis* (22.6 ± SEM 8.3%), were detected with no significant variation across honey samples (Two-way ANOVA, Sidak’s Multiple comparison tests). Only *Frischella perrara* was significantly enriched in honey harvested in the spring. Additional symbiotic bacteria were also detected with limited variation across honey samples (Sup. figure 11c), with the exception of 5 species (*Enterobacter sp. SA187*, *Klebsiella oxytoca*, *Leuconostoc pseudomesenteroides*, *Sodalis glossinidius* and *Sodalis praecaptivus*), which were significantly more abundant in autumn honey samples (Figure 5c). To further explore the potential role of such seasonal associations between bacteria and honeybee, we functionally annotated the reads aligning against these 5 genomes specifically and identified significantly enriched functions for each species (Figure 5d). For example, *Enterobacter sp. SA187* and *Leuconostoc pseudomesenteroides* were both enriched in metabolic functions related with carbohydrate process and glycogenesis, while *Sodalis glossinidius* and *Sodalis praecaptivus* were enriched in functions related to transposition and amino-acid transfer, respectively.

In conclusion, these observations revealed that metagenomic analyses of honey samples derived from a static apiary allow for the non-invasive characterisation of honeybee pathogens and core microbiome, but can also describe specific adaptations of the microbiota, in particular for processing of season-specific sugars and, more surprisingly, for functions associated with genome remodelling.

### 3.6 The Direct-shotgun metagenomic approach quantified the number of natural varroa falls more accurately

The original hives of the studied honey samples showed relatively low contamination by varroa mites at harvest time (Sup. figure 1). Nevertheless, an average of 47 ± 17 reads corresponding to varroa DNA were detected across the sequenced libraries and we sought to investigate the relationship between the abundances detected by shotgun metagenomics with the natural falls of varroa over periods of 2 weeks, 1 month and 2 months preceding honey collection. As a validation step, we also correlated these abundances with the abundances obtained after re-alignment of each sequenced library specifically against the *Varroa destructor* genome.

Overall, the Direct-SM approach correlated better with the average of natural varroa falls over a one-month period (R= 0.68), while the SM approach inversely correlated with all measurements of varroa natural fall (Figure 6). Interestingly, these correlations with *V. destructor* genomic alignments showed similar trends pointing to this result not being due to pipeline classification or normalization steps, but, most likely, due to the different DNA extraction methods that were employed. Therefore, the Direct-SM approach appears to be more suitable for the biomonitoring of monthly varroa infestation.

**Figure 6:**
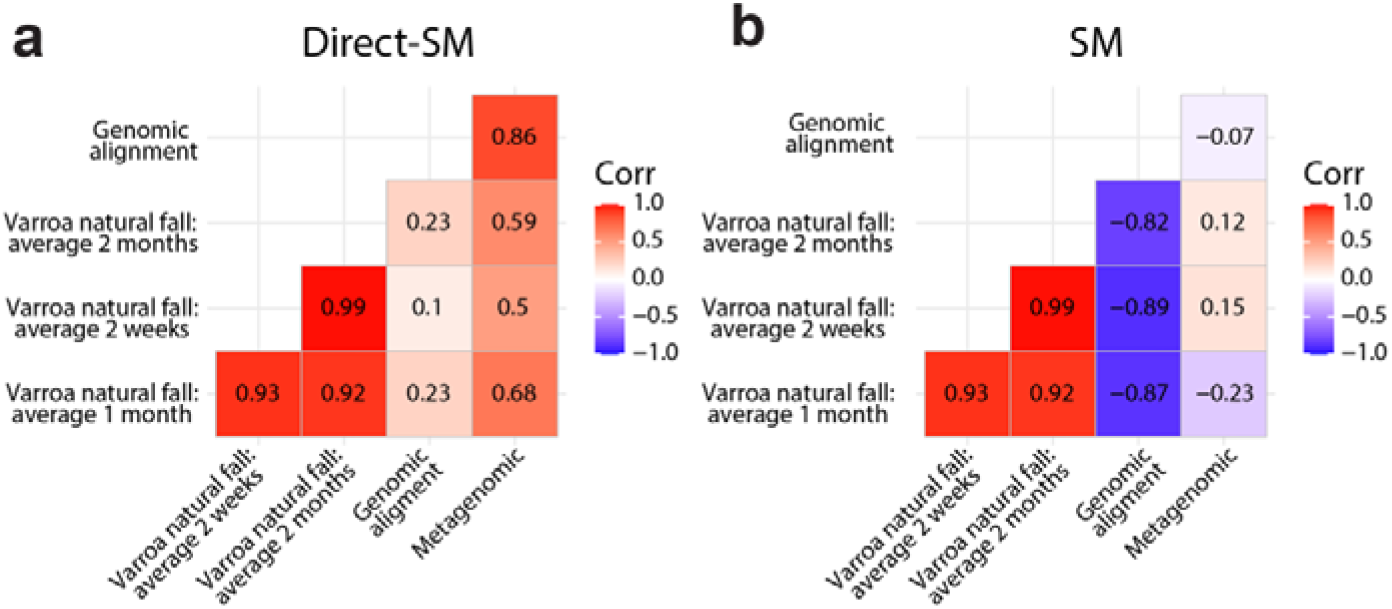
The Direct-shotgun metagenomic methodology correlates better with the number of natural varroa falls. Correlation matrices were built by comparing counts per million from genomic alignment against the *Varroa destructor* genome (‘Genomic alignment’) with the average of varroa natural falls per day over periods of 2 weeks, 1 month, and 2 months (‘Varroa natural fall’) and with *Varroa* species abundance detected by our metagenomic pipeline (‘Metagenomic’), comparing (a) Direct-SM and (b) SM approaches.

## 4. DISCUSSION

The assessment of the complex ecological niche that honeybees occupy has only recently begun to be examined through the use of shotgun metagenomics (*47*–*49*). In this study we have developed a direct approach for shotgun metagenomics (Direct-SM) and a bioinformatic pipeline to assess the species composition of honey samples. Our analyses revealed that Direct-SM offers an integrative approach to molecularly dissect not only the seasonal adaptive behaviours of foragers, but also their gut microbiota composition and hive pests, such as varroa.

It is important to note that shotgun metagenomics studies often suffer from high rates of false positives and are rarely either computationally or analytically validated. We have optimised and validated our metagenomic classification pipeline using simulated samples that resemble the species composition of actual honey samples. We have also optimised the DNA extraction step to reduce biased enrichment for certain species. For example, some DNA extraction methods may be more efficient in lysing Gram-positive bacteria than other methods (*66*, *67*). In fact, we show that standard methods of DNA extraction may fail to record hive varroa infestation compared with the shotgun metagenomic extraction methodology described herein.

Bees directly depend on plants to meet their energetic and nutrient needs. Importantly, pollen itself is known to facilitate associative learning in honeybees (*68*), and pollen quality is associated with cognitive function (*13*), emphasising the importance of accurately determining honeybee foraging patterns. We show that Direct-SM reported the presence in honey of similar families as those detected by melissopalynological analyses. In particular, the detection of tree species (such as pine and eucalyptus) with high abundance is consistent with the findings of previous studies describing the foraging patterns of urban and suburban apiaries (*36*–*38*). The identification of plants through next generation sequencing still remains a challenge due to the relative lack of sequenced plant genomes (*61*). Nevertheless, it can be used in addition to melissopalynological analysis, in particular for quantifying foraging behaviours associated with non-pollen harvests.

Our methodology is further suitable for describing species interactions in the honeybee gut. Our study reports for the first time the detection and seasonal stability of core and non-core gut bacterial species in honey through honey shotgun metagenomics. Notably, *Frischella perrara* was more abundantly detected in spring, which is in accordance with the findings of previous studies (*42*). Interestingly, autumn honey was enriched in *Sodalis* species, which are known to be more abundant in solitary bee species (*69*). This may characterise bee transition to overwintering, as honeybees begin their non-productive and less active period.

Functional annotation of the autumn-specific bacteria identified functions related to carbohydrate metabolism, amino acid process and transport, and transposition. Roles for bacteria in carbohydrate metabolism have been described in the past and are crucial for polysaccharide breakdown of pollen and nectar sugars (*70*). Likewise, amino acid transport appears to occur frequently between the gut microbiome and honeybees (*71*). *Sodalis glossinidius* was enriched in functions related to transposition, consistent with a recent study that evaluated the genomic landscape of *Sodalis glossonidius* and showed that a large part of it encoded for transposition-related pseudogenes (*72*). Importantly, *Sodalis praecaptivus* is considered to be the only currently known free-living form of *Sodalis* bacteria (*73*), while *Sodalis glossonidius* is an insect endosymbiont (*74*). The exact function of these transposition-related pseudogenes has not yet been identified, but owing to their identification here in autumn-specific microbiota, it may be worth considering how transposition is related to host colonisation and how both transposition and associated metabolic functions relate to honeybee overwintering. On the other hand, endosymbionts are considered to have enabled insects to adapt in new environments. Recently, they have also been found to provide various functions to their insect hosts, including pesticide detoxification (*75*), and to communicate with the insect immune system, providing among others increased protection against other bacterial and viral species (*76*).

As the honeybee gut receives increasingly more attention, it is important to develop non-invasive methods that are able to capture the diversity of the “shared gut” within the colony. The constant exchange of honey and beebread between honeybees allows them to transfer gut microbes between generations, potentially contributing to hive-specific microbiomes and behavioural variation. Non-invasive methods, such as direct shotgun metagenomics, can complement existing studies of single-honeybee 16S rDNA in order to sample colony microbiomes.

The methodologies described here are aimed at constituting a field-deployable system allowing for the description of the complexity of the honeybee ecological niche in a non-invasive and precise manner. This will enable not only the assessment of the environmental diversity of habitats surrounding apiaries during specific times of the year, but also the evaluation of honeybee health in an integrative manner taking into account food type, gut microbiome composition and beehive pest invasion.

## Acknowledgements

This work was funded by a Marie Skłodowska-Curie Individual Fellowship (798082) to S.P. and by a Greek Ministry of Rural Development and Food grant (program for production and marketing of beekeeping) to E.S. G.A.P. was supported by the Operational Program Competitiveness, Entrepreneurship and Innovation, NSRF 2014-2020, Action code: MIS 5002562, co-financed by Greece and the European Union (European Regional Development Fund). We would like to thank Charlotta Wiren for field assistance, Nikolaos Kalavros for assistance in bioinformatic analyses, and Foteini Ravini and Anastasios Kampolis from the CheMa Laboratories.

## Data Accessibility Statement

The datasets have been deposited in the Sequence Read Archive (SRA) from NCBI under the BioProject <u>ID PRJNA735927</u>. Source codes are available in Github at https://github.com/AGalanis97/Direct-shotgun-metagenomics-pub.

## Author Contributions

A.G., P. H., S.P. designed the research. A.G., P.V., M.R., G.A.P., S.P. acquired the data. A.G., V.H., conducted wet lab. A.G., S.P., analysed the data. A.G., M.R., developed code for data analysis. A.G., S.P. provided data visualisation. A.G., P.V., M.R. provided data validation. A.G., S.P. wrote the original draft. G.A.P., P.H., S.P. reviewed and edited manuscript. E.M.C.S., S.P. supervised the study. G.A.P., E.M.C.S., S.P. provided funds.

## References

1. A. M. Klein, V. Boreux, F. Fornoff, A. C. Mupepele, G. Pufal, Relevance of wild and managed bees for human well-being. Curr. Opin. Insect Sci. 26, 82–88 (2018).

2. V. Patel, N. Pauli, E. Biggs, L. Barbour, B. Boruff, Why bees are critical for achieving sustainable development. Ambio, 49–59 (2020).

3. J. Belsky, N. K. Joshi, Impact of biotic and abiotic stressors on managed and feral bees. Insects. 10 (2019).

4. D. Goulson et al., Bee declines driven by combined stress from parasites, pesticides, and lack of flowers. Science. 347, 1255957 (2015).

5. K. M. Smith et al., Pathogens, pests, and economics: Drivers of honey bee colony declines and losses. Ecohealth. 10, 434–445 (2013).

6. S. Klein, A. Cabirol, J.-M. Devaud, A. B. Barron, M. Lihoreau, Why Bees Are So Vulnerable to Environmental Stressors. Trends Ecol Evol (2016).

7. M. A. Aizen, L. D. Harder, The Global Stock of Domesticated Honey Bees Is Growing Slower Than Agricultural Demand for Pollination. Curr. Biol. 19, 915–918 (2009).

8. W. Glenny et al., Honey bee (Apis mellifera) colony health and pathogen composition in migratory beekeeping operations involved in California almond pollination. PLoS One. 12, 1–24 (2017).

9. M. Simone-Finstrom et al., Migratory management and environmental conditions affect lifespan and oxidative stress in honey bees. Sci. Rep. 6, 1–10 (2016).

10. T. L. Dynes, J. A. Berry, K. S. Delaplane, B. J. Brosi, J. C. De Roode, Reduced density and visually complex apiaries reduce parasite load and promote honey production and overwintering survival in honey bees. PLoS One. 14, 1–16 (2019).

11. L. Castelli et al., Impact of Nutritional Stress on Honeybee Gut Microbiota, Immunity, and Nosema ceranae Infection. Microb. Ecol. 80, 908–919 (2020).

12. B. Branchiccela et al., Impact of nutritional stress on the honeybee colony health. Sci. Rep. 9, 1–11 (2019).

13. G. Di Pasquale et al., Variations in the availability of pollen resources affect honey bee health. PLoS One. 11, 1–15 (2016).

14. D. Naug, Nutritional stress due to habitat loss may explain recent honeybee colony collapses. Biol. Conserv. 142, 2369–2372 (2009).

15. J. Liberti, P. Engel, The gut microbiota - brain axis of insects. Curr. Opin. Insect Sci. (2020).

16. E. V. S. Motta, K. Raymann, N. A. Moran, Glyphosate perturbs the gut microbiota of honey bees. Proc. Natl. Acad. Sci. U. S. A. 115, 10305–10310 (2018).

17. J. C. Jones et al., Gut microbiota composition is associated with environmental landscape in honey bees. Ecol. Evol. 8, 441–451 (2018).

18. L. Paris et al., Honeybee gut microbiota dysbiosis in pesticide/parasite co-exposures is mainly induced by Nosema ceranae. J. Invertebr. Pathol. 172 (2020).

19. L. Zhu, S. Qi, X. Xue, X. Niu, L. Wu, Nitenpyram disturbs gut microbiota and influences metabolic homeostasis and immunity in honey bee (Apis mellifera L.). Environ. Pollut. 258 (2020)..

20. J. T. Kerr et al., Climate change impacts on bumblebees converge across continents. Science. 349, 177–180 (2015).

21. M. Kammerer, S. C. Goslee, M. R. Douglas, J. F. Tooker, C. M. Grozinger, Wild bees as winners and losers: Relative impacts of landscape composition, quality, and climate. Glob. Chang. Biol. 27, 1250–1265 (2021).

22. M. Matzrafi, Climate change exacerbates pest damage through reduced pesticide efficacy. Pest Manag. Sci. 75, 9–13 (2019).

23. I. Delcour, P. Spanoghe, Literature review: Impact of climate change on pesticide use. Food Res. Int. 68, 7–15 (2015).

24. S. G. Potts et al., Safeguarding pollinators and their values to human well-being. Nature. 540, 220–229 (2016).

25. J. Settele, J. Bishop, S. G. Potts, Climate change impacts on pollination. Nat. Plants. 2, 1–3 (2016).

26. M. M. López-Uribe, V. A. Ricigliano, M. Simone-Finstrom, Defining Pollinator Health: A Holistic Approach Based on Ecological, Genetic, and Physiological Factors. Annu. Rev. Anim. Biosci. 8, 269–294 (2020).

27. C. J. Perry, E. Søvik, M. R. Myerscough, A. B. Barron, Rapid behavioral maturation accelerates failure of stressed honey bee colonies. Proc. Natl. Acad. Sci. U. S. A. 112, 3427–3432 (2015).

28. M. L. Smith et al., The dominant axes of lifetime behavioral variation in honey bees. bioRxiv, 1–28 (2021).

29. European Food Safety Authority, Towards an integrated environmental risk assessment of multiple stressors on bees: review of research projects in Europe, knowledge gaps and recommendations. EFSA J. 12 (2014).

30. G. Gilioli, G. Sperandio, F. Hatjina, A. Simonetto, Towards the development of an index for the holistic assessment of the health status of a honey bee colony. Ecol. Indic. 101, 341–347 (2019).

31. K. Deiner et al., Environmental DNA metabarcoding: Transforming how we survey animal and plant communities. Mol. Ecol. 26, 5872–5895 (2017).

32. M. F. Breed et al., The potential of genomics for restoring ecosystems and biodiversity. Nat. Rev. Genet. 20, 615–628 (2019).

33. I. Kafantaris, G. D. Amoutzias, D. Mossialos, Foodomics in bee product research: a systematic literature review. Eur. Food Res. Technol. 247, 309–331 (2021).

34. J. Hawkins et al., Using DNA Metabarcoding to Identify the Floral Composition of Honey: A New Tool for Investigating Honey Bee Foraging Preferences. PLoS One. 10, e0134735 (2015).

35. N. Peel et al., Semi-quantitative characterisation of mixed pollen samples using MinION sequencing and Reverse Metagenomics (RevMet). Methods Ecol. Evol. 2019, 1–12 (2019).

36. A. E. Samuelson, R. J. Gill, E. Leadbeater, Urbanisation is associated with reduced Nosema sp. infection, higher colony strength and higher richness of foraged pollen in honeybees. Apidologie. 51, 746–762 (2020).

37. N. De Vere et al., Using DNA metabarcoding to investigate honey bee foraging reveals limited flower use despite high floral availability. Sci. Rep. 7 (2017).

38. D. B. Sponsler, D. Shump, R. T. Richardson, C. M. Grozinger, Characterizing the floral resources of a North American metropolis using a honey bee foraging assay. Ecosphre. 11 (2020).

39. L. Jones et al., Shifts in honeybee foraging reveal historical changes in floral resources. Commun. Biol. 4, 1–10 (2021).

40. P. Engel, V. G. Martinson, N. A. Moran, Functional diversity within the simple gut microbiota of the honey bee. Proc. Natl. Acad. Sci. U. S. A. 109, 11002–11007 (2012).

41. K. M. Ellegaard, P. Engel, Genomic diversity landscape of the honey bee gut microbiota. Nat. Commun. 10 (2019), doi:10.1038/s41467-019-08303-0.

42. L. Kešnerová, O. Emery, M. Troilo, B. Erkosar, P. Engel, Gut microbiota structure differs between honey bees in winter and summer. ISME J. (2019), doi:10.1101/703512.

43. J. H. Yun, M. J. Jung, P. S. Kim, J. W. Bae, Social status shapes the bacterial and fungal gut communities of the honey bee. Sci. Rep. 8, 1–11 (2018).

44. R. S. Cornman et al., Pathogen Webs in Collapsing Honey Bee Colonies. PLoS One. 7, e43562 (2012).

45. P. D’Alvise, V. Seeburger, K. Gihring, M. Kieboom, M. Hasselmann, Seasonal dynamics and co-occurrence patterns of honey bee pathogens revealed by high-throughput RT-qPCR analysis. Ecol. Evol. 9, 10241–10252 (2019).

46. G. Bonilla-Rosso, T. Steiner, F. Wichmann, E. Bexkens, P. Engel, Honey bees harbor a diverse gut virome engaging in nested strain-level interactions with the microbiota. Proc. Natl. Acad. Sci. U. S. A. (2020).

47. S. Bovo et al., Shotgun metagenomics of honey DNA: Evaluation of a methodological approach to describe a multi-kingdom honey bee derived environmental DNA signature. PLoS One. 13 (2018).

48. S. Bovo, V. J. Utzeri, A. Ribani, R. Cabbri, L. Fontanesi, Shotgun sequencing of honey DNA can describe honey bee derived environmental signatures and the honey bee hologenome complexity. Sci. Rep. 10, 1–17 (2020).

49. H. Wirta, N. Abrego, K. Miller, T. Roslin, E. Vesterinen, DNA traces the origin of honey by identifying plants, bacteria and fungi. Sci. Rep. 11, 1–14 (2021).

50. K. S. Delaplane, J. van der Steen, E. Guzman-Novoa, Standard methods for estimating strength parameters of Apis mellifera colonies. J. Apic. Res. 52, 1–12 (2013).

51. A. Shcherbina, FASTQSim: Platform-independent data characterization and in silico read generation for NGS datasets. BMC Res. Notes. 7, 1–12 (2014).

52. V. R. Marcelino et al., CCMetagen: comprehensive and accurate identification of eukaryotes and prokaryotes in metagenomic data. Genome Biol. 21:103 (2020).

53. B. Buchfink, C. Xie, D. H. Huson, Fast and sensitive protein alignment using DIAMOND. Nat. Methods. 12, 59–60 (2015).

54. D. E. Wood, J. Lu, B. Langmead, Improved metagenomic analysis with Kraken 2. Genome Biol. 20, 257 (2019).

55. K. P. Keegan, E. M. Glass, F. Meyer, F. Martin, S. Uroz, Eds. (Springer New York, New York, NY, 2016.

56. H. Li, Minimap2: pairwise alignment for nucleotide sequences. Bioinformatics. 34, 3094–3100 (2018).

57. M. I. Love, W. Huber, S. Anders, Moderated estimation of fold change and dispersion for RNA-seq data with DESeq2. Genome Biol. 15, 1–21 (2014).

58. M. B. Pereira, M. Wallroth, V. Jonsson, E. Kristiansson, Comparison of normalization methods for the analysis of metagenomic gene abundance data. BMC Genomics. 19, 1–17 (2018).

59. L. Pantano, DEGreport: Report of DEG analysis. (2019), doi:10.18129/B9.bioc.DEGreport.

60. A. Kassambara, F. Mundt, Package Factoextra: Visualization of a Correlation MatrixExtract and Visualize the Results of Multivariate Data Analyses (2019).

61. B. S. Maitner et al., The bien r package: A tool to access the Botanical Information and Ecology Network (BIEN) database. Methods Ecol. Evol. 9, 373–379 (2018).

62. E. A. Franzosa et al., Species-level functional profiling of metagenomes and metatranscriptomes. Nat. Methods. 15 (2018).

63. D. Kim, J. M. Paggi, C. Park, C. Bennett, S. L. Salzberg, Graph-based genome alignment and genotyping with HISAT2 and HISAT-genotype. Nat. Biotechnol. 37, 907–915 (2019).

64. C. A. Kwadha, G. O. Ong’Amo, P. N. Ndegwa, S. K. Raina, A. T. Fombong, The biology and control of the greater wax moth, Galleria mellonella. Insects. 8, 1–17 (2017).

65. R. S. Schwarz et al., Honey bee colonies act as reservoirs for two Spiroplasma facultative symbionts and incur complex, multiyear infection dynamics. Microbiologyopen. 3, 341–355 (2014).

66. K. O. Bøifot, J. Gohli, L. V. Moen, M. Dybwad, Performance evaluation of a new custom, multi-component DNA isolation method optimized for use in shotgun metagenomic sequencing-based aerosol microbiome research. Environ. Microbiomes. 15, 1–23 (2020).

67. F. Yang et al., Assessment of fecal DNA extraction protocols for metagenomic studies. Gigascience. 9, 1–12 (2020).

68. F. Muth, D. R. Papaj, A. S. Leonard, Bees remember flowers for more than one reason: Pollen mediates associative learning. Anim. Behav. 111, 93–100 (2016).

69. B. E. R. Rubin, J. G. Sanders, K. M. Turner, N. E. Pierce, S. D. Kocher, Social behaviour in bees influences the abundance of *Sodalis* (Enterobacteriaceae) symbionts. R. Soc. Open Sci. 5, 180369 (2018).

70. H. Zheng et al., Division of labor in honey bee gut microbiota for plant polysaccharide digestion. Proc. Natl. Acad. Sci. U. S. A. 116, 25909–25916 (2019).

71. H. Zheng, J. E. Powell, M. I. Steele, C. Dietrich, N. A. Moran, Honeybee gut microbiota promotes host weight gain via bacterial metabolism and hormonal signaling. Proc. Natl. Acad. Sci. U. S. A. 114, 4775–4780 (2017).

72. I. Goodhead et al., Large-scale and significant expression from pseudogenes in Sodalis glossinidius – a facultative bacterial endosymbiont. Microb. Genomics. 6 (2020).

73. A. Chari et al., Phenotypic characterization of sodalis praecaptivus sp. nov., a close non-insect-associated member of the sodalis-allied lineage of insect endosymbionts. Int. J. Syst. Evol. Microbiol. 65, 1400–1405 (2015).

74. A. K. Snyder, C. M. Mcmillen, P. Wallenhorst, R. V. M. Rio, The phylogeny of Sodalis-like symbionts as reconstructed using surface-encoding loci. FEMS Microbiol. Lett. 317, 143–151 (2011).

75. Q. Su, X. Zhou, Y. Zhang, Symbiont-mediated functions in insect hosts. Commun. Integr. Biol. 6 (2013).

76. L. Eleftherianos, J. Atri, J. Accetta, J. C. Castillo, Endosymbiotic bacteria in insects: Guardians of the immune system? Front. Physiol. 4 MAR, 1–10 (2013).

