## Supplementary information for "Bee foraging preferences, microbiota and pathogens revealed by direct shotgun metagenomics of honey"

**Supplementary Figures**

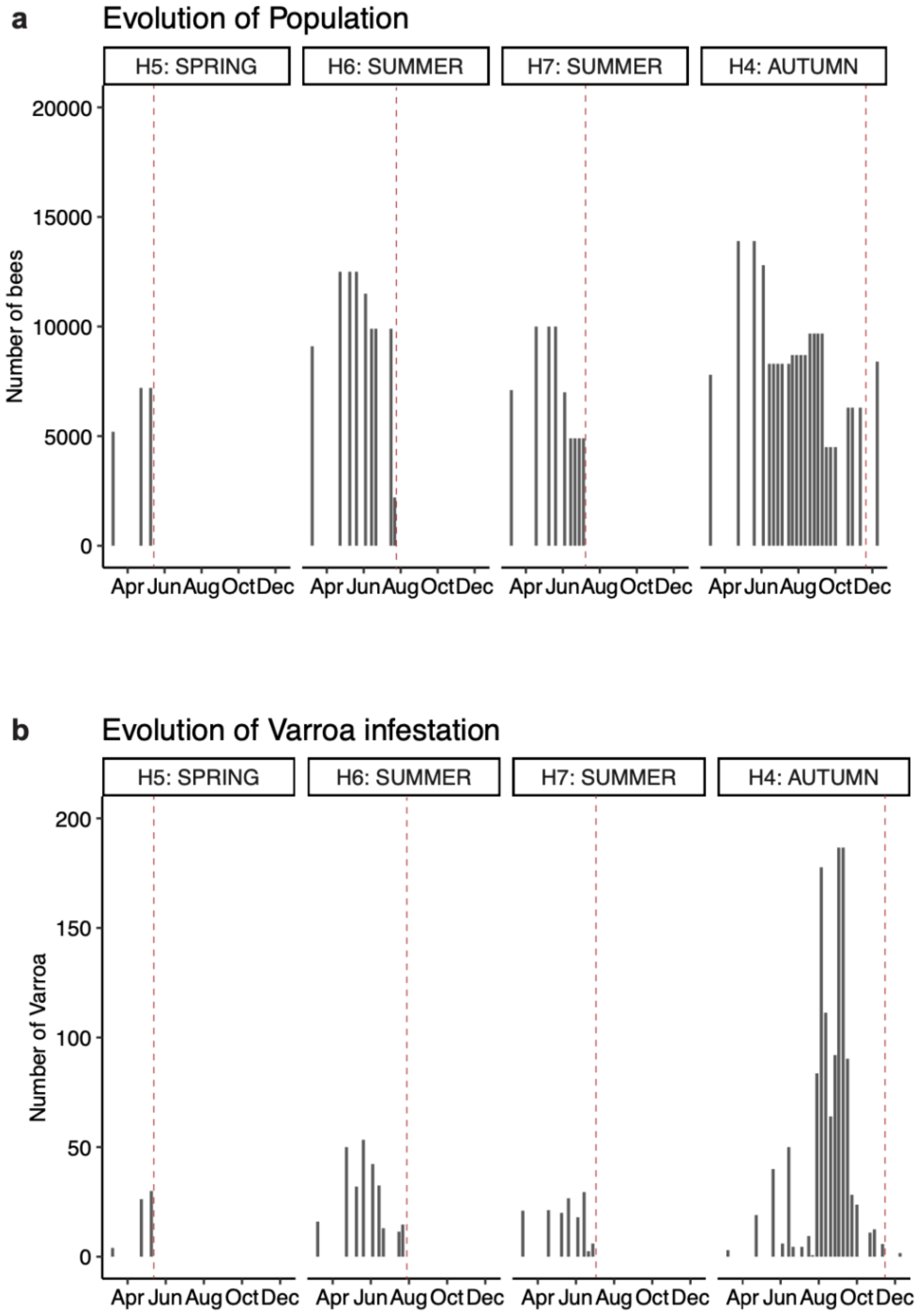


**Supplementary Figure 1: Apiary monitoring.**

History of the honeybee population (a) and of varroa infestation (b) at the time of honey sampling (red line).

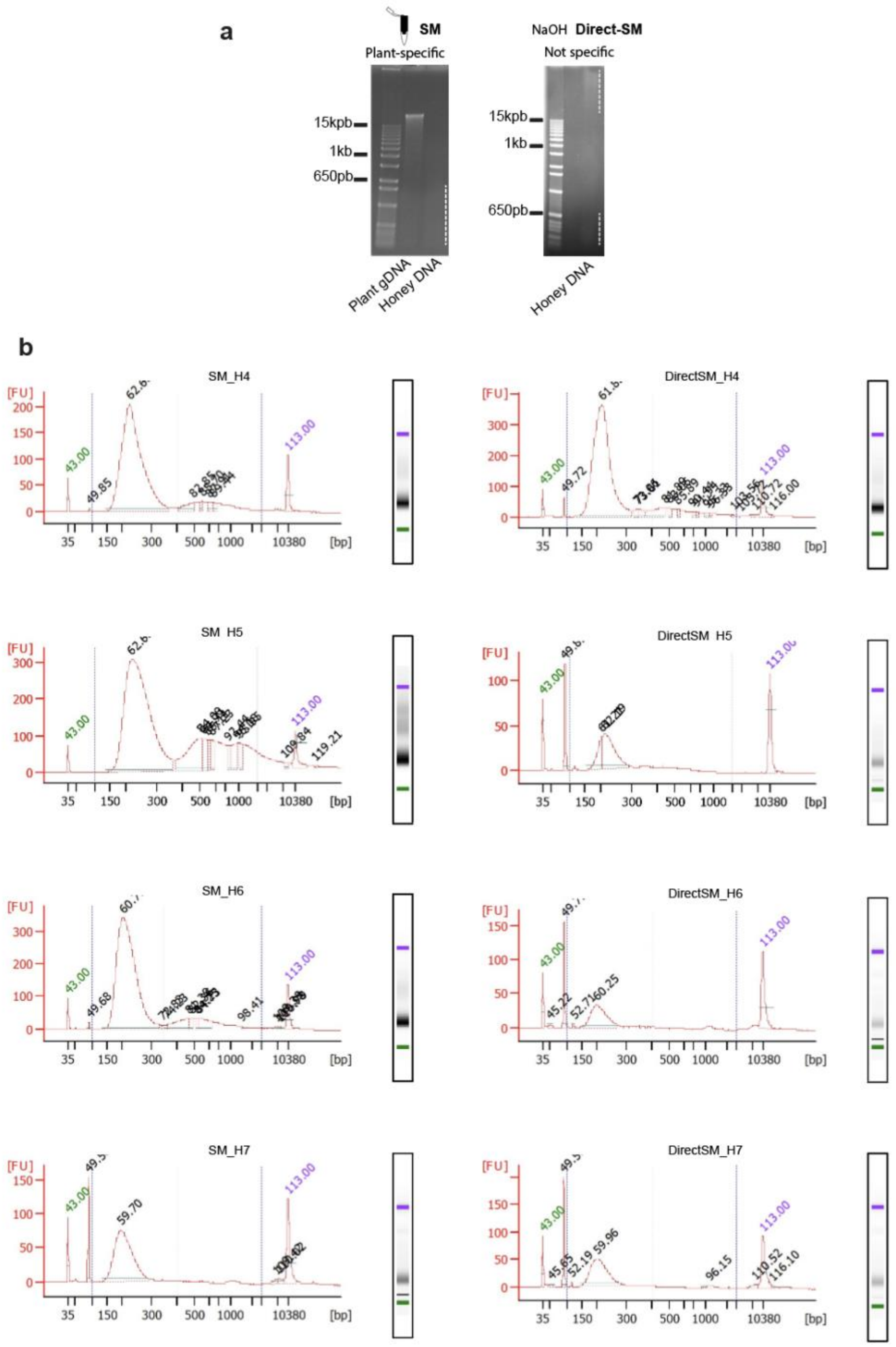


**Supplementary Figure 2: Quality controls of the DNA extraction and sequenced libraries.**

(a) Agarose gel after DNA extraction using respectively SM and Direct-SM methodology. Dash lines indicated degraded DNA from honey samples. (b) Bioanalyzer of the 8 libraries prior sequencing.

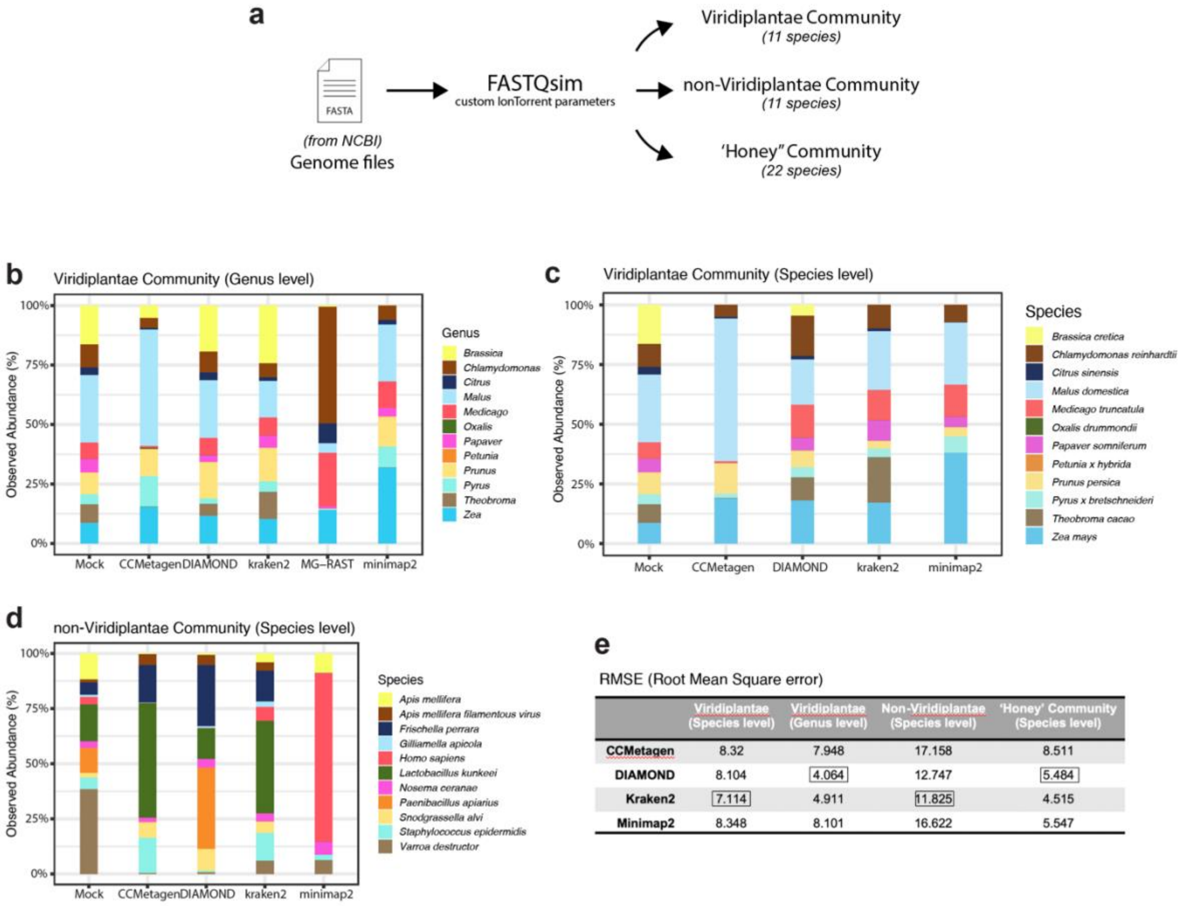


| | Viridiplantae (Species level) | Viridiplantae (Genus level) | Non-Viridiplantae (Species level) | 'Honey' Community (Species level) |
|---|---|---|---|---|
| **CCMetagen** | 8.32 | 7.948 | 17.158 | 8.511 |
| **DIAMOND** | 8.104 | 4.064 | 12.747 | 5.484 |
| **Kraken2** | 7.114 | 4.911 | 11.825 | 4.515 |
| **Minimap2** | 8.348 | 8.101 | 16.622 | 5.547 |

**Supplementary Figure 3: Evaluation of the taxonomic classifiers and genomic aligners.**

(a) Flowchart depicting the details bioinformatics analysis leading to construction of 3 main mock communities. (b - d) Barplots showing the abundance distribution according to the build communities. (e) Summary table of the mean square error (RMSE) associated to each taxonomic classifier with the mock build communities.

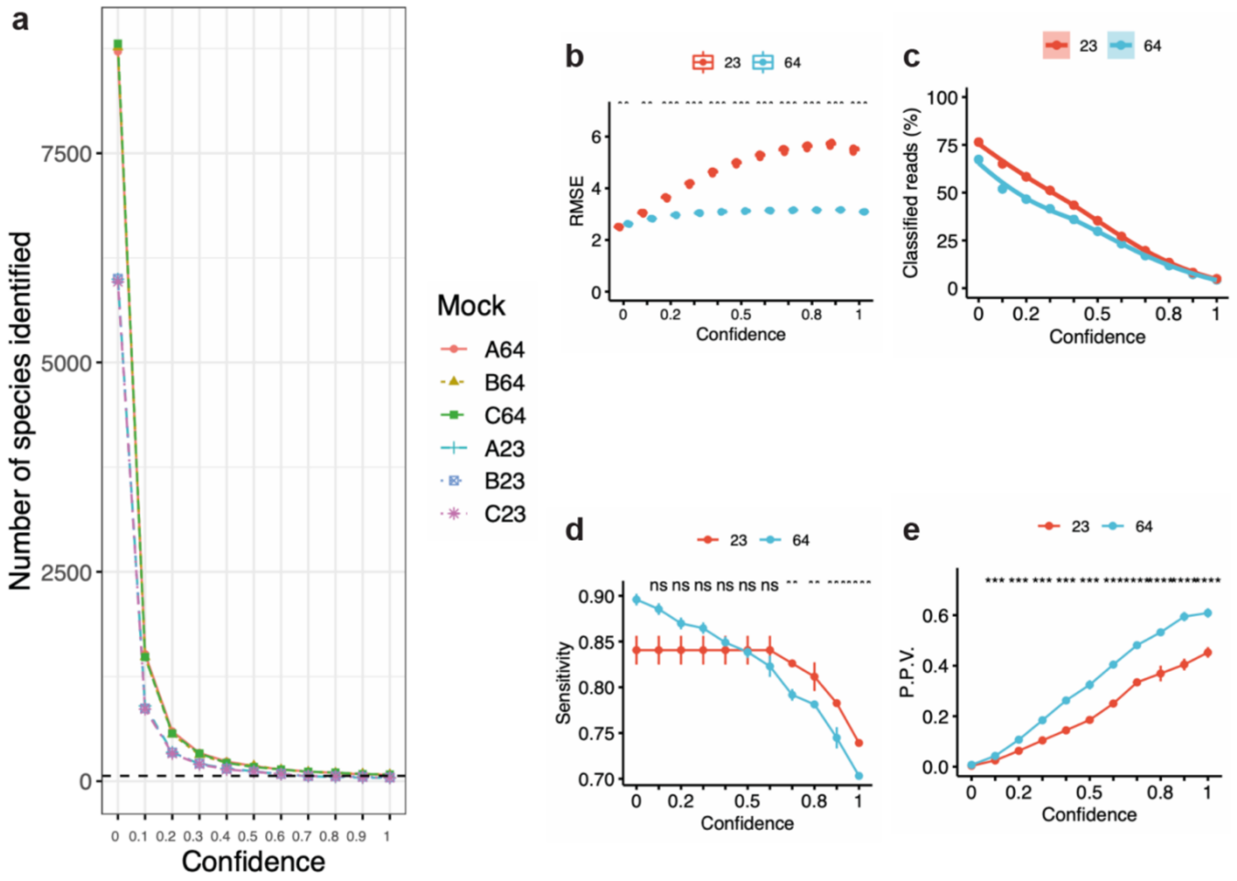


**Supplementary Figure 4: Evaluation of Kraken 2 confidence threshold in relation with species content.**

(a) absolute number of species content, (b) Root Mean Square error (RMSE), (c) Percentage of classified reads, (d) Sensitivity, (e) Positive Predictive Value (P.P.V).

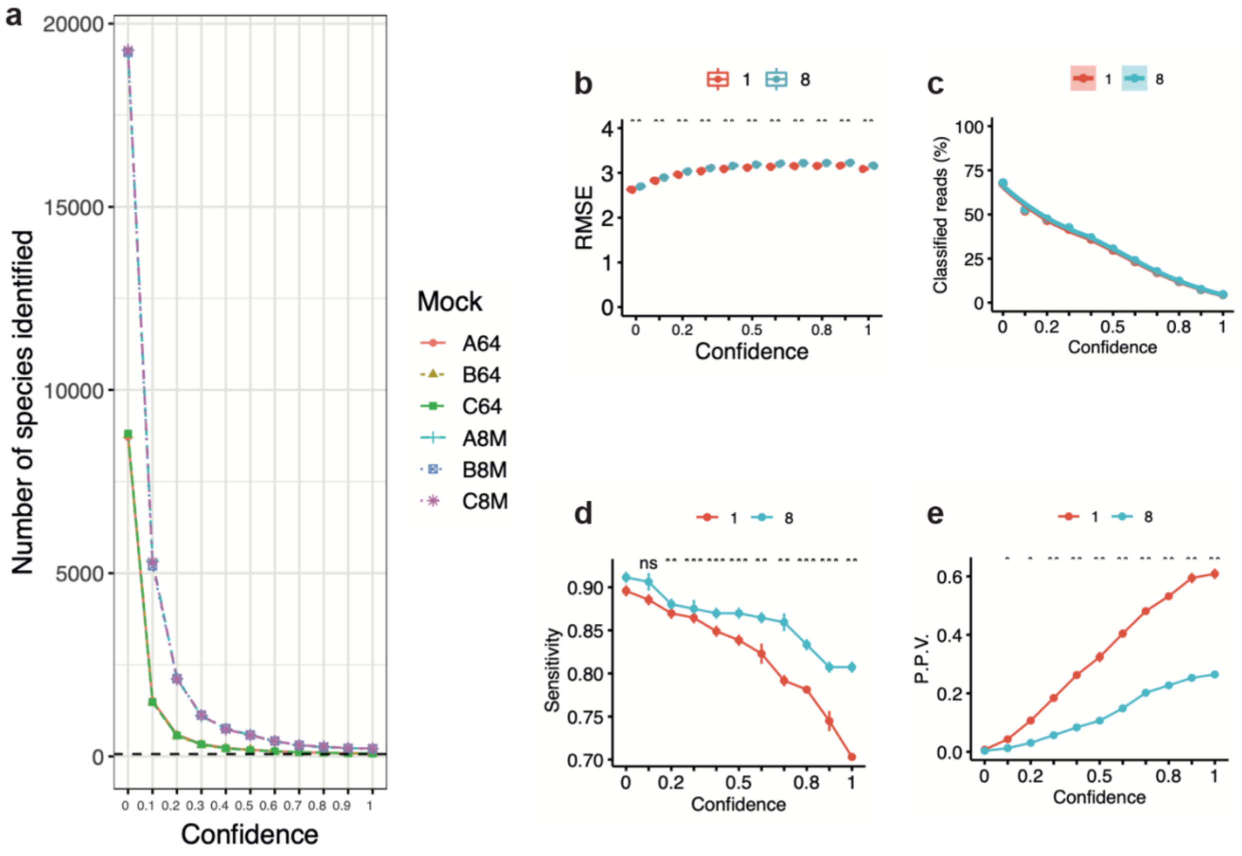


**Supplementary Figure 5: Evaluation of Kraken 2 confidence threshold in relation with Library size.**

(a) absolute number of species content, (b) Root Mean Square error (RMSE), (c) Percentage of classified reads, (d) Sensitivity, (e) Positive Predictive Value (P.P.V).

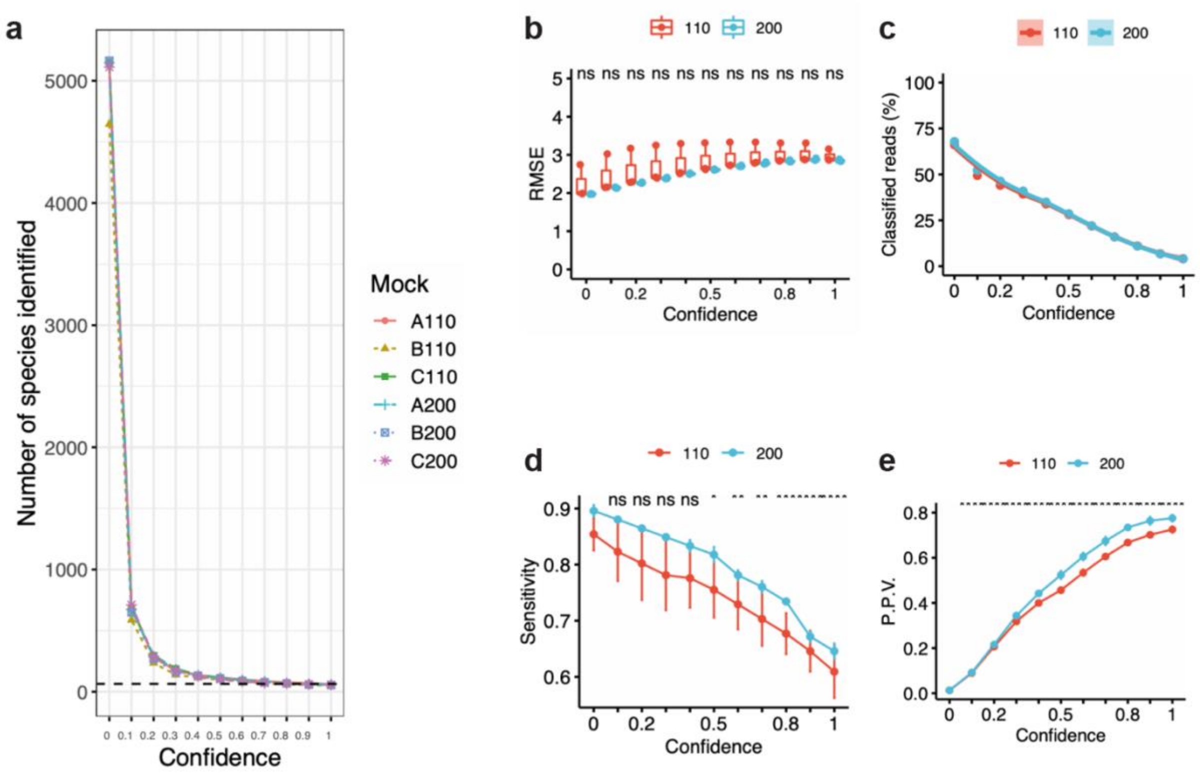


**Supplementary Figure 6: Evaluation of Kraken 2 confidence threshold in relation with read length.**

(a) absolute number of species content, (b) Root Mean Square error (RMSE), (c) Percentage of classified reads, (d) Sensitivity, (e) Positive Predictive Value (P.P.V).

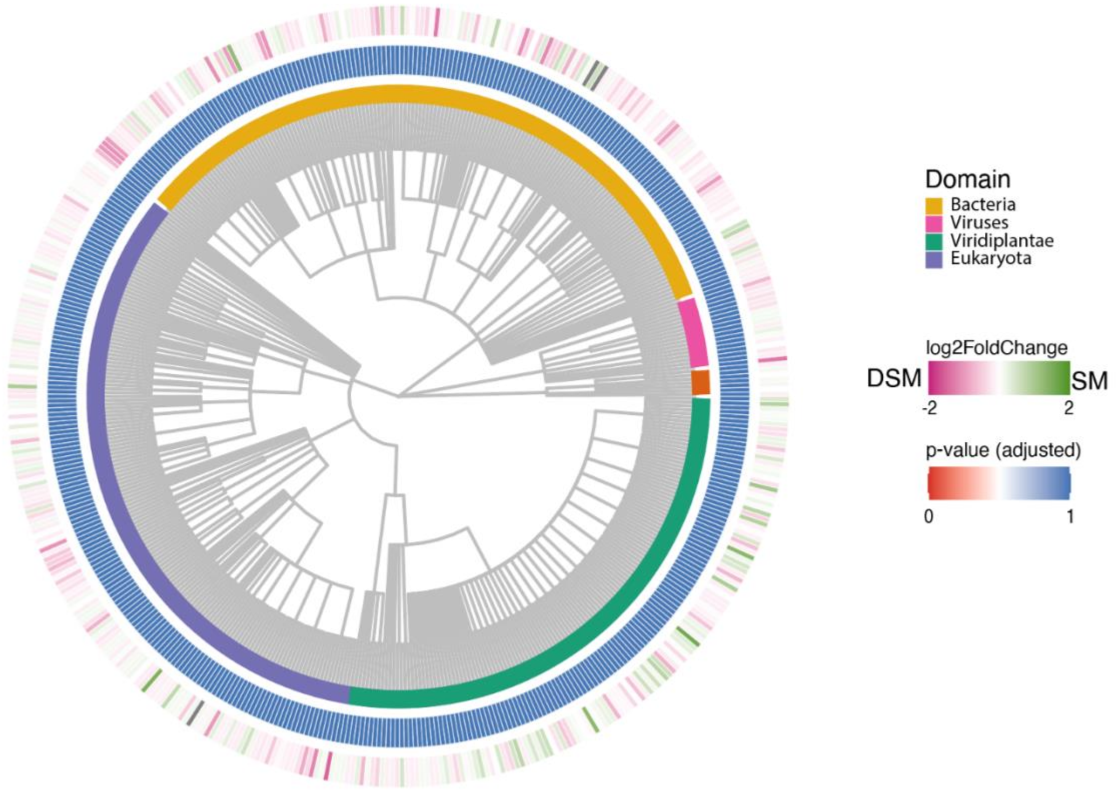


**Supplementary Figure 7: Summary of family distribution and abundance variation between methodologies.**

Phylogenetic distribution of the 477 families identified by honey metagenomic and classified by domains. Log2FoldChange indicate the direction of change towards greater abundance between the classic SM methodology (green, 4 technical replicates) and the direct-SM methodology (pink, 4 technical replicates). The adjusted p-values were calculated using DESeq2 Wald test with Benjamini and Hocheberg multiple comparisons.

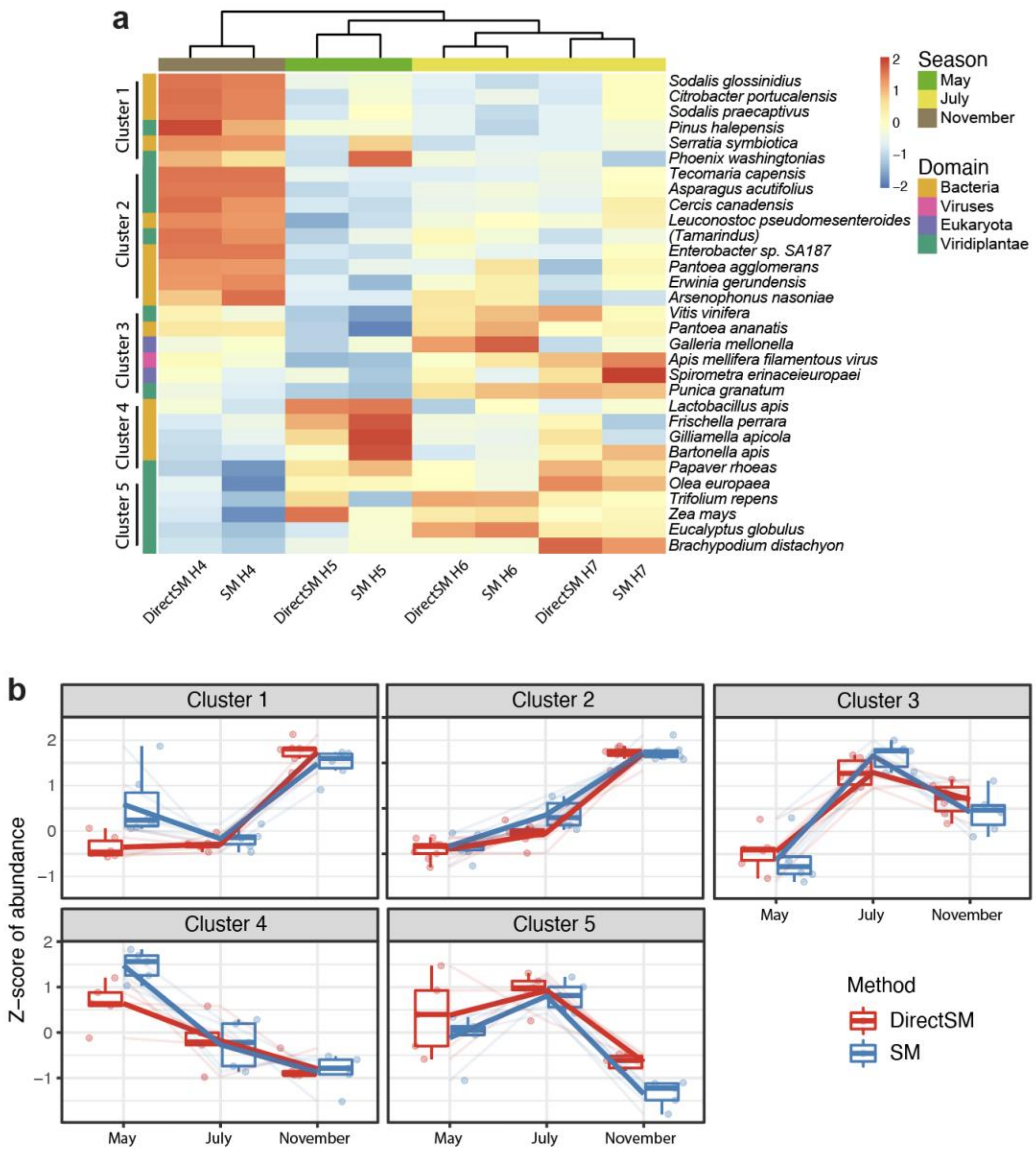


**Supplementary Figure 8: Heatmap and hierarchical clusters.**

(a) Heatmap depicting hierarchical clustering of the 31 species identified to be differentially abundant across the 8 sequenced libraries (filtered for species with a minimum coverage of 50 reads). Side annotation show the groups of species with similar dynamic of abundance across seasons. (b) Detail of abundance dynamic across the hierarchical clusters associated with a specific season (May: spring, July: summer, November: autumn).

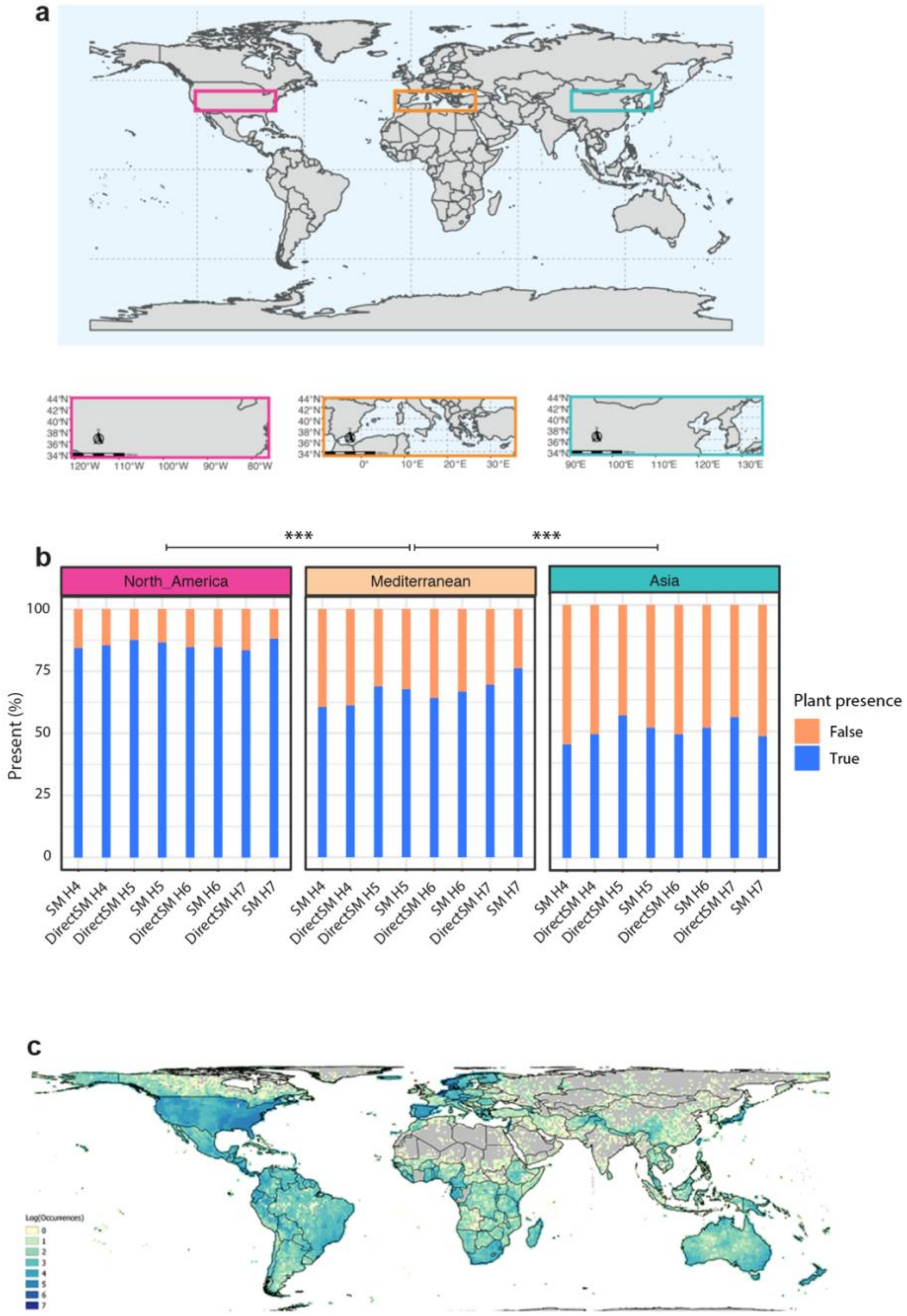


**Supplementary Figure 9: Plant validation (*in silico*).**

(a) World map showing the 3 equivalent areas used to calculate the number of plant occurrences in our study compared to the plant occurrences associated with each of these areas. (b) Results of plant occurrence for each analyzed area. The statistical test used is the Fisher 'exact test (p value <0.0001). (c) World map of the plant occurrence from our study across the entire world. Map generated by https://bien.nceas.ucsb.edu/bien/biendata/

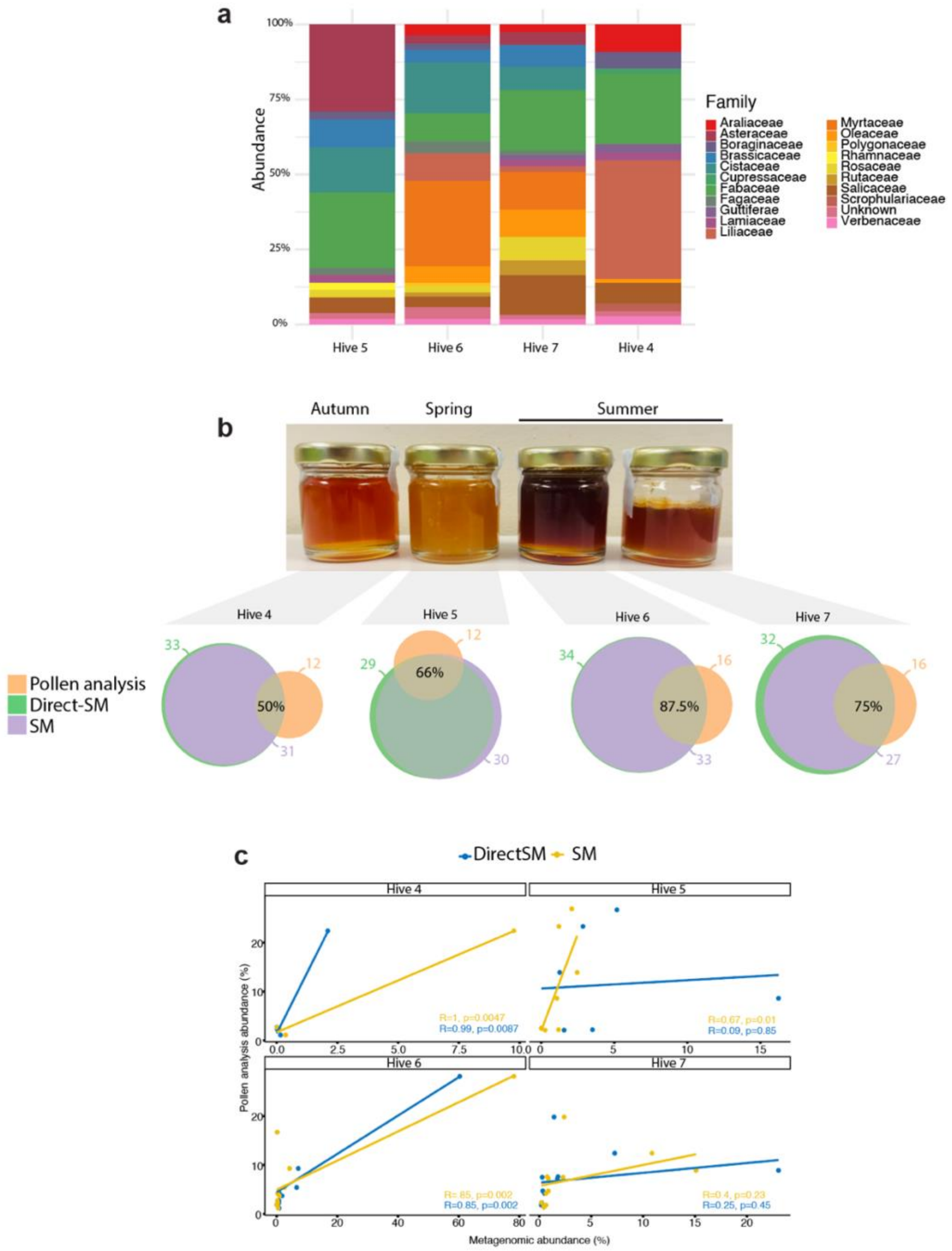


**Supplementary Figure 10: Plant validation (pollen analysis).**

(a) Barplots showing the distribution of the 20 pollen families identified by melissopalynology. (b) Upper panel: colour of the 4 honeys collected in this study across 3 seasons. Lower panel: Euler plots showing the superposition of the families of plants identified for each of the 2 metagenomic analyses (Direct-SM and SM) with the one identified by pollen analysis. The number of families associated with each technic is indicated as well as the % of families overlapping the pollen analysis. (c) Correlation plots of family plant abundances between the 2 metagenomic analyses and the pollen analysis for each hive. The correlation coefficient and significance are indicated.

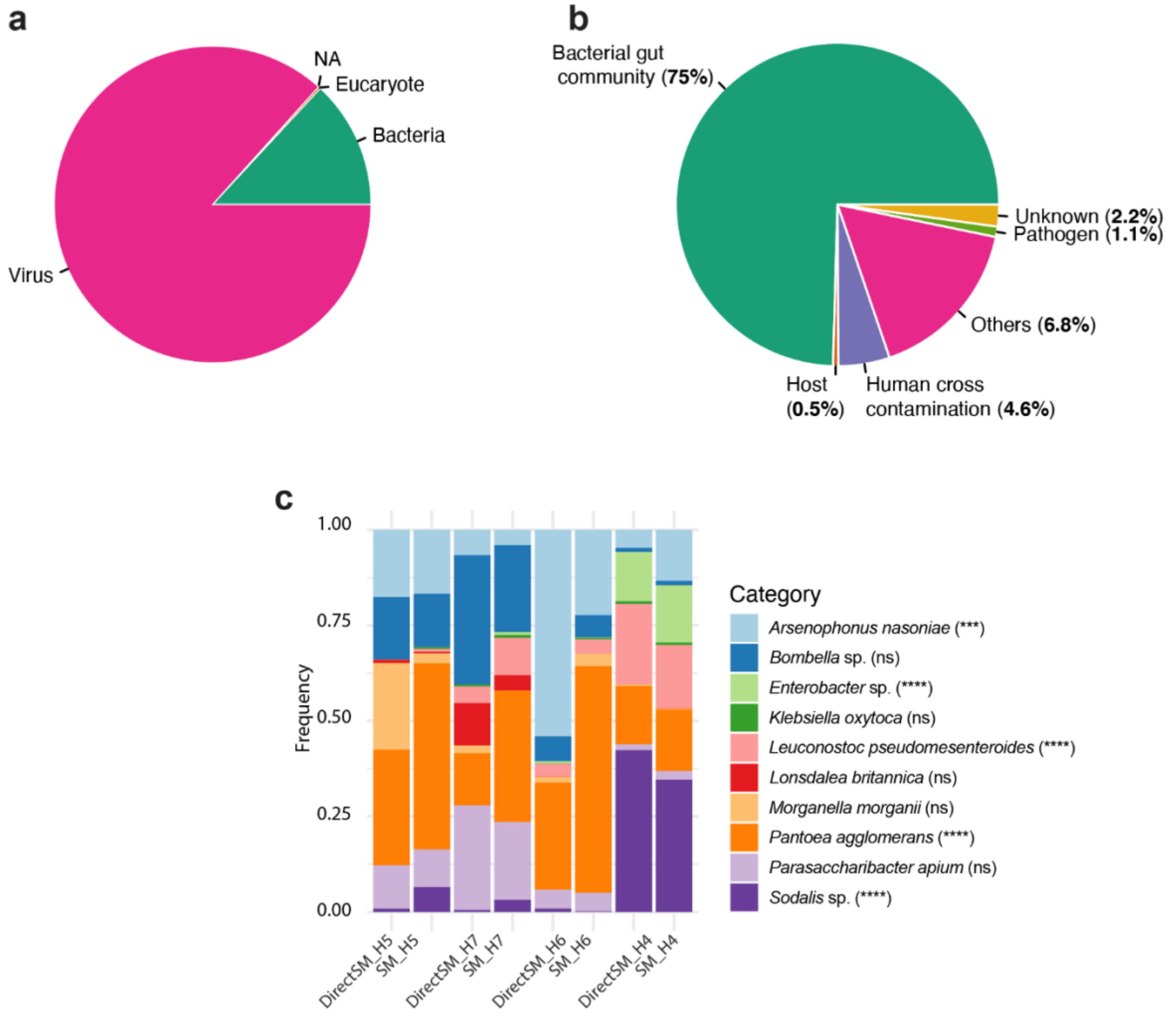


**Supplementary Figure 11: Non-plant distribution and association with honeybee**

Pies depicting the proportion of DNA constituting all the sequenced reads from non-plant species (a) without filtering and (b) after filtering the *Apis meliferra* filamentous virus and species categorization according to their relationship with the honeybees. Only species with more than 100 reads are shown (Table S3). (c) Barplots of the symbiotic bacteria distribution not belonging to bacteria previously described as core or noncore. Species with significant variations are annotated with stars.

**Supplementary Tables**

**Table S1: Viridiplantea validation**

| **Family** | **Genus** | **Identified species (Metagenomic)** | **Abundance %** | **Season specific *** | **Identified species (Visual)** | **Common name (Greek name)** | **Validation level** |
|---|---|---|---|---|---|---|---|
| Asparagaceae | *Asparagus* | *A. officinalis* | 2,90 | Autumn | *A. Acutifolius* | Wild asparagus (Αγριο σπαράγγι) | Genus |
| Malvaceae | *Bombax* | *B. ceiba* | 0,06 | N/A | None | Red cotton tree | **Not Validated** (Asia) |
| Poaceae | *Brachypodium* | *B. distachyon* | 0,36 | Summer | *B. distachyon* | Purple false brome | Species |
| Fabaceae | *Castanospermum* | *C. australe* | 0,08 | Autumn | None (Ornamental) | Moreton Bay chestnut | **Not Validated** |
| Fabaceae | *Ceratonia* | *C. siliqua* | 0,15 | Autumn | *C. siliqua* | Carob (Χαρουπιά) | Species |
| Fabaceae | *Cercis* | *C. canadensis* | 0,18 | Autumn | *C. siliquastrum* (Endemic), *C. canadensis* (Ornamental) | Judas tree or forest pansy (Κουτσουπιά) | Species |
| Cucurbitaceae | *Citrullus* | *C. lanatus* | 0,07 | N/A | *C. lanatus* | Watermelon (Καρπούζι) | Species |
| Rutaceae | *Citrus* | *C. sinensis* | 0,30 | N/A | *C. aurantium* | Orange Tree (Πορτοκαλιά) | Genus |
| Arecaceae | *Cocos* | *C. nucifera* | 0,11 | N/A | None | Coconut Tree (Καρύδα) | **Not Validated** (Tropical) |
| Cucurbitaceae | *Ecballium* | *E. elaterium* | 0,08 | N/A | *E. elaterium* | Squirting cucumber (Πικραγγουριά / Γαιδαγγουριά) | Species |
| Phrymaceae | *Erythranthe* | *E. guttata* | 0,09 | N/A | *E. guttata* (Ornamental) | Monkeyflower (Μιλουλος) | Species |
| Myrtaceae | *Eucalyptus* | *E. grandis* | 12,48 | Summer | *E. globulus* | Eucalyptus (Ευκάλυπτος) | Genus |

| | | | | | | | |
|---|---|---|---|---|---|---|---|
| Asteraceae | *Glebionis* | *G. coronaria* | 0,09 | Spring | *G. coronaria* | Crown daisy (Χρυσάνθεμο / Μαντηλίδα) | Species |
| Fabaceae | *Libidibia* | *L. coriaria* | 0,08 | Autumn | None | Guaracabuya | **Not Validated** (Tropical) |
| Oleaceae | *Ligustrum* | *L. quihoui* | 0,06 | N/A | *L. japonicum, L. vulgaris* | Privet (Λιγουστρος) | Genus |
| Magnoliaceae | *Liriodendron* | *L. tulipifera* | 0,06 | N/A | *L. tulipifera aureomarginatum* (Ornamental) | Tulip Tree (Λιριόδεντρο) | Species |
| Fabaceae | *Lotus* | *L. japonicus* | 0,06 | N/A | *L. cytisoides* | Grey Birdsfoot Trefoil (Λωτός ο κυτισοειδής) | Genus |
| Rosaceae | *Malus* | *M. domestica* | 0,09 | N/A | *M. domestica* | Appletree (Μηλιά) | Species |
| Fabaceae | *Medicago* | *M. truncatula* | 0,19 | N/A | *M. orobrea* | Alfalfa (Μηδίκη) | Genus |
| Moraceae | *Morus* | *M. alba* | 0,15 | N/A | *M. alba* | White mulberry (Λευκή μουριά) | Species |
| Oleaceae | *Olea* | *O. europaea* | 1,96 | Summer | *O. europaea* | Olive Tree (Ελιά) | Species |
| Orobanchaceae | *Orobanche* | *O. crenata* | 0,11 | N/A | *O. crenata* | Broomrape (Οροβάγχη) | Species |
| Papaveraceae | *Papaver* | *P. rhoeas, P. somniferum* | 13,98 | Spring | *P. rhoeas* | Poppy (Παπαρούνα) | Species |
| Arecaceae | *Phoenix* | *P. dactylifera* | 12,50 | Autumn | *P. washingtonias* | Date palm (χουρμαδια) | Genus |
| Pinaceae | *Pinus* | *P. pinea, P. pinaster, P. taeda, P. roxburghii* | 47,42 | Autumn | *P. halepensis* | Pine tree (Πεύκο) | Genus |
| Anacardiaceae | *Pistacia* | *P. vera* | 0,44 | N/A | *P. lentescus* | Pistachio (Φυσιτκιά) | Genus |
| Rosaceae | *Prunus* | *P. dulcis* | 0,05 | N/A | *P. dulcis* | Almond Tree (Αμυγδαλιά) | Genus |
| Lythraceae | *Punica* | *P. granatum* | 0,35 | Summer | *P. granatum* | Pomegranate (Ροδιά) | Species |
| Fagaceae | *Quercus* | *Q. suber* | 0,10 | N/A | *Q. coccifera, Q. Aegilops* | Kermes oak (Πουρνάρι) | Genus |

| | | | | | | | |
|---|---|---|---|---|---|---|---|
| Brassicaceae | *Raphanus* | *R. sativus* | 0,35 | N/A | *R. raphanistrum* | Wild radish (Ραπανίδα) | Genus |
| Rosaceae | *Rosa* | *R. chinensis* | 0,15 | N/A | *R. chinensis* (Ornamental) | China Rose (Τριαντάφυλλο) | Species |
| Lamiaceae | *Salvia* | *S. miltiorrhiza* | 0,07 | N/A | *S. officinalis* | Sage (Φασκόμηλο) | Genus |
| Schisandraceae | *Schisandra* | *S. sphenanthera* | 0,05 | N/A | None (Ornamental) | Magnolia vine | **Not Validated** (Asia) |
| Fabaceae | *Tamarindus* | *T. indica* | 0,13 | Autumn | None | Tamarind | **Not Validated** (Tropical) |
| Bignoniaceae | *Tecomaria* | *T. capensis* | 0,25 | Autumn | *T. capensis* | Red Cape Honeysuckle (Βιγνόνια αειθαλής) | Species |
| Lamiaceae | *Thymbra* | *T. capitata* | 0,07 | Summer | *T. capitatus* | Conehead thyme (Κεφαλωτό Θυμάρι) | Species |
| Fabaceae | *Trifolium* | *T.repens* | 0,63 | Summer | Various *Trifolium* spp. | (Wild) Clover (τριφυλλι (αγριοτρίφυλλο) | Species |
| Poaceae | *Triticum* | *T. aestivum* | 0,09 | N/A | *T. aegilops, T. comosa* | Wheat (Σταρι) | Genus |
| Vitaceae | *Vitis* | *V. vinifera* | 1,22 | Summer | *V. vinifera subs. Silverstris* | Vine (Αμπελος) | Species |
| Sapindaceae | *Xanthoceras* | *X. sorbifolium* | 0,05 | N/A | None | Yellowhorn | **Not Validated (Tropical)** |
| Poaceae | *Zea* | *Z. mays* | 2,41 | Summer | *Z. mays* | Corn (Καλαμποκι) | Species |

* The specificity associated with a season was assessed by hierarchical clustering analysis (See methods).

3

**Table S2: Main inconsistences between pollen and metagenomic analyses**

| Family | Identified species | Common name | Detected in Hive: |
|---|---|---|---|
| *<u>Never found in Pollen analysis but detected >1% in both SM and Direct-SM metagenomic analysis</u>* | | | |
| Arecaceae | *Phoenix washingtonias* | Date palm | H4, H6, H7 |
| Asparagaceae | *Asparagus Acutifolius* | Wild asparagus | H4 |
| Lythraceae | *Punica granatum* | Pomegranate | H7 |
| Papaveraceae | *Papaver rhoeas* | Poppy | H5, H6, H7 |
| Pinaceae | *Pinus halepensis* | Pine tree | H4, H5 |
| Poaceae | *Brachypodium distachyon* | Purple false brome | H6, H7 |
| Vitaceae | *Vitis vinifera subs. Silverstris* | Vine | H6, H7 |
| *<u>Never found in Metagenomic analysis but detected >1% in pollen analysis</u>* | | | |
| Araliaceae* | *Hedera helix* | Ivy | H4, H6, H7 |
| Boraginaceae* | | | H4, H6, H7 |
| Scrophulariaceae* | | | H4 |

* No family or close family sequenced genome

6 **Table S3: Non-plant species classification and text mining references**

| Species | NCBI search in abstract content: ‘bee’/’honey’/’Apis’ with 'Species name' | Relation with *Apis mellifera* | Total reads | Category |
|---|---|---|---|---|
| *Acinetobacter baumannii* | https://pubmed.ncbi.nlm.nih.gov/?term=24883072+28855917+29138732+30596703+29904274+30108574+31186997+31516358+26408816 | N/A - False positive (human pathogen) | 229 | Human cross contamination |
| *Acinetobacter johnsonii* | [ ] | N/A - False positive (human pathogen) | 290 | Human cross contamination |
| *Acinetobacter pittii* | [ ] | N/A - False positive (in human skin) | 553 | Human cross contamination |
| *Lactobacillus apinorum* | Not apply | Gut Microbiota | 222 | Bacterial gut community |
| *Lactobacillus kunkeei* | https://pubmed.ncbi.nlm.nih.gov/?term=22427985+24466119+24516438+23991051+24740297+25792062+24944337+25953738+25768309+24148670+25768110+25319366+25301653+26140264+27324340+27114887+28572315+28144419+28243545+28346815+28291793+28663705+29472795+29386588+29321884+28966892+28934240+28717593+29038770+27140688+29125851+30353179+30990837+30223435+30249054+29914555+30100619+30379893+30683856+31285515+31243442+31378758+32075309+32275718+32222941+32443465+31664160+31739261+31800608+31027777+30269313+29648528+29264966+27776911+27633178+26121394+25609652+25239902+25211052+24478297+22659204 | Gut Microbiota | 286241 | Bacterial gut community |
| *Apis cerana* | Not apply | Host | 142 | Host |
| *Apis mellifera* | Not apply | Host | 2150 | Host |

| | | | | |
|---|---|---|---|---|
| *Arsenophonus nasoniae* | https://www.ncbi.nlm.nih.gov/pmc/articles/PMC4941583/ | Symbiotic | 1820 | Bacterial gut community |
| *Bartonella apis* | https://pubmed.ncbi.nlm.nih.gov/?term=21594072+28234349+28291793+28435856+29232373+29386588+29321884+28966892+27140688+29608282+30353179+30990837+29764668+29914555+30100619+30249635+30683856+31703071+31753958+32275718+30467713+27730366+26537852 | Core - Gut Microbiota | 1651 | Bacterial gut community |
| *Bombella sp. ESL0368* | [ ] | Gut Microbiota | 598 | Bacterial gut community |
| *Citrobacter portucalensis* | https://pubmed.ncbi.nlm.nih.gov/18683661/ | Gut microbiota - pathogen in relation with hibernation | 1063 | Pathogen |
| *Cutibacterium acnes* | https://pubmed.ncbi.nlm.nih.gov/?term=23802986+25597924+27280049+26580653+28204822+29138732+29247241+27187328+30596703+30132547+29401750+28534835+31252651+32012913+31613567+25872535+25215662+25062791+24496237+24063779+17196372 | N/A - False positive (in human skin) | 2996 | Human cross contamination |
| *Cutibacterium granulosum* | [ ] | N/A - False positive (in human skin) | 216 | Human cross contamination |
| *Delftia tsuruhatensis* | [ ] | N/A - False positive (human pathogen) | 453 | Human cross contamination |
| *Enterobacter cancerogenus* | [ ] | N/A - False positive (human pathogen) | 1999 | Human cross contamination |
| *Enterobacter hormaechei* | [ ] | N/A - False positive (human pathogen) | 107 | Human cross contamination |
| *Enterobacter sp. SA187* | https://pubmed.ncbi.nlm.nih.gov/18683661/ | Gut microbiota - symbiotic in relation with hibernation | 1965 | Bacterial gut community |
| *Erwinia gerundensis* | [ ] | N/A - plant pathogen | 280 | Others |

| | | | | |
|---|---|---|---|---|
| *Frischella perrara* | https://pubmed.ncbi.nlm.nih.gov/?term=24358254+24740297+25991680+25768309+25874551+25210772+25852743+26140264+26330094+27118586+27803186+28291793+28386455+27720045+29164717+29232373+29386588+29321884+29876054+28698604+28715431+27140688+29125851+30846803+30353179+29764668+29914555+30100619+30249635+30379893+30683856+31271530+31624548+31703071+23606484+31800608+28207182+27717118+25239900 | Core - Gut microbiota - seasonal, caste variability | 8110 | Bacterial gut community |
| *Galleria mellonella* | Not apply | Pathogen | 2404 | Pathogen |
| *Gilliamella apicola* | https://pubmed.ncbi.nlm.nih.gov/?term=22558460+22829932+23111871+24740297+25991680+25768309+25768110+25148082+25874551+25880915+25319366+25395644+25210772+25852743+26140264+26330094+26011669+26623177+26437644+27803186+28346815+28291793+28386455+28435856+29164717+29232373+29386588+29321884+28966892+29876054+28698604+27140688+29125851+29608282+29635372+30353179+30990837+29764668+30575755+29914555+30100619+30249635+30379893+31271530+31243442+31540209+31378758+32005933+32275718+23041637+23606484+30467713+29624166+29397399+25805518+25239900+25211052+25053814+23347062+23060052+22307297 | Core - Gut microbiota - seasonal, caste variability | 2248 | Bacterial gut community |
| *Habropoda laboriosa* | [ ] | Unknown | 353 | Unknown |
| *Herbaspirillum huttiense* | [ ] | N/A - False positive (human pathogen) | 110 | Human cross contamination |

| | | | | |
|---|---|---|---|---|
| *Hydrogenophilus thermoluteolus* | [ ] | Unknown | 164 | Unknown |
| *Klebsiella oxytoca* | https://pubmed.ncbi.nlm.nih.gov/18683661/ | Gut microbiota - symbiotic in relation with hibernation | 108 | Bacterial gut community |
| *Lactobacillus apis* | Not apply | Core - Firm5 | 214 | Bacterial gut community |
| *Lactobacillus delbrueckii* | Not apply | Core - Firm5 | 219 | Bacterial gut community |
| *Lactobacillus kullabergensis* | Not apply | Core - Firm5 | 145 | Bacterial gut community |
| *Lactobacillus sp. Fhon2N* | Not apply | Core - Firm5 | 321 | Bacterial gut community |
| *Lactococcus lactis* | https://pubmed.ncbi.nlm.nih.gov/?term=19156201+22615573+23028506+24587122+25953738+24936378+25880915+27114887+28912763+25884548+30249054+31271530+30092318+25211052+18286279 | N/A - False positive (food contamination) | 2524 | Human cross contamination |
| *Leuconostoc pseudomesenteroides* | http://amsdottorato.unibo.it/8626/1/Ph.D_Thesis_Alberoni_D.pdf | N/A - Found in Gut of Osmia | 2790 | Bacterial gut community |
| *Liquorilactobacillus nagelii* | [ ] | Unknown | 119 | Unknown |
| *Malassezia globosa* | [ ] | N/A - False positive (in human skin) | 297 | Human cross contamination |
| *Malassezia restricta* | [ ] | N/A - False positive (in human skin) | 1036 | Human cross contamination |
| *Moraxella osloensis* | [ ] | N/A - slug pathogen | 210 | Others |
| *Morganella morganii* | https://pubmed.ncbi.nlm.nih.gov/18683661/ | Gut microbiota - symbiotic in relation with hibernation | 144 | Bacterial gut community |
| *Apis mellifera filamentous virus* | https://pubmed.ncbi.nlm.nih.gov/26008705/ | Weak pathogen | 2699492 | Pathogen |
| *Bacteriophage sp.* | https://pubmed.ncbi.nlm.nih.gov/32341166/ | Unknown | 854 | Unknown |

| | | | | |
|---|---|---|---|---|
| *eukaryotic synthetic construct* | Not apply | N/A - GO | 168 | Others |
| *Myoviridae sp.* | [ ] | Unknown | 824 | Unknown |
| *Siphoviridae sp.* | [ ] | Unknown | 5970 | Unknown |
| *uncultured Bacteroidetes bacterium* | Not apply | Unknown | 240 | Unknown |
| *Pantoea agglomerans* | https://pubmed.ncbi.nlm.nih.gov/?term=26330094+27279628+28663705+30379893+32107443+25895542+18683661 | Gut Microbiota | 3280 | Bacterial gut community |
| *Pantoea ananatis* | https://pubmed.ncbi.nlm.nih.gov/?term=29799863 | N/A - plant pathogen | 1197 | Others |
| *Parasaccharibacter apium* | https://pubmed.ncbi.nlm.nih.gov/26875068/ | Symbiotic | 636 | Bacterial gut community |
| *Pseudomonas oryzihabitans* | Not apply | N/A - False positive (human pathogen) | 2755 | Human cross contamination |
| *Pseudomonas syringae* | Not apply | N/A - False positive (human pathogen) | 399 | Human cross contamination |
| *Pseudomonas viridiflava* | Not apply | N/A - False positive (human pathogen) | 190 | Human cross contamination |
| *Serratia symbiotica* | https://pubmed.ncbi.nlm.nih.gov/32518251/ | N/A - Aphid symbion | 68376 | Others |
| *Sodalis glossinidius* | https://pubmed.ncbi.nlm.nih.gov/?term=30109092 | Gut microbiota - symbiotic in relation with sociality | 1357 | Bacterial gut community |
| *Sodalis praecaptivus* | https://pubmed.ncbi.nlm.nih.gov/?term=30109092 | Gut microbiota - symbiotic in relation with sociality | 4110 | Bacterial gut community |
| *Spirometra erinaceieuropaei* | [ ] | Unknown | 768 | Unknown |
| *Spiroplasma melliferum* | https://pubmed.ncbi.nlm.nih.gov/24771723/ | Pathogen | 886 | Pathogen |
| *Staphylococcus capitis* | https://pubmed.ncbi.nlm.nih.gov/?term=29657429+31762646 | N/A - False positive (in human skin) | 2926 | Human cross contamination |

| | | | | |
|---|---|---|---|---|
| *Staphylococcus epidermidis* | Not apply | N/A - False positive (in human skin) | 677 | Human cross contamination |
| *Staphylococcus hominis* | Not apply | N/A - False positive (in human skin) | 158 | Human cross contamination |
| *Streptococcus gordonii* | https://pubmed.ncbi.nlm.nih.gov/?term=26330195+28392803+30026515 | N/A - False positive (in human cavital teeth) | 193 | Human cross contamination |
| *Streptococcus sanguinis* | Not apply | N/A - False positive (in human skin) | 120 | Human cross contamination |
| *Streptococcus thermophilus* | Not apply | N/A - False positive (in human skin) | 364 | Human cross contamination |
| *Tatumella ptyseos* | [ ] | N/A - False positive (human pathogen) | 838 | Human cross contamination |
| *Tatumella sp. TA1* | Not apply | N/A - False positive (human pathogen) | 2179 | Bacterial gut community |
| *Zymobacter palmae* | [ ] | N/A - False positive (human pathogen) | 103 | Human cross contamination |

7
